# Ranked Subtree Prune and Regraft

**DOI:** 10.1101/2023.05.16.541039

**Authors:** Lena Collienne, Chris Whidden, Alex Gavryushkin

## Abstract

Phylogenetic trees are a mathematical formalisation of evolutionary histories between organisms, species, genes, cancer cells, etc. For many applications, e.g. when analysing virus transmission trees or cancer evolution, (phylogenetic) time trees are of interest, where branch lengths represent times. Computational methods for reconstructing time trees from (typically molecular) sequence data, for example Bayesian phylogenetic inference using Markov Chain Monte Carlo (MCMC) methods, rely on algorithms that sample the treespace. They employ tree rearrangement operations such as SPR (Subtree Prune and Regraft) and NNI (Nearest Neighbour Interchange) or, in the case of time tree inference, versions of these that take times of internal nodes into account. While the classic SPR tree rearrangement is well-studied, its variants for time trees are less understood, limiting comparative analysis for time tree methods.

In this paper we consider a modification of the classical SPR rearrangement on the space of ranked phylogenetic trees, which are trees equipped with a ranking of all internal nodes. This modification results in two novel treespaces, which we propose to study. We begin this study by discussing algorithmic properties of these treespaces, focusing on those relating to the complexity of computing distances under the ranked SPR operations as well as similarities and differences to known tree rearrangement based treespaces. Surprisingly, we show the counterintuitive result that adding leaves to trees can actually decrease their ranked SPR distance, which may have an impact on the results of time tree sampling algorithms given uncertain “rogue taxa”.

## 1 Introduction

Phylogenetic trees are used to display evolutionary relationships, for example between organisms, species, or genes, and are usually inferred from DNA, RNA, amino acid, or other types of sequence data. Typical applications include reconstructing the evolutionary history of a set of species, e.g. to construct the tree of life [24], analysing transmission patterns of viruses [30, 34], and investigating cancer evolution [3]. The goal is the same in all applications: finding a phylogenetic tree that best explains evolutionary relationship between given (sequence) data, represented by the leaves of the tree.

Many methods for inferring phylogenetic trees from sequence data use tree sampling or treespace search algorithms. These include Bayesian methods, for example the software packages BEAST [14], BEAST2 [7], MrBayes [23], and RevBayes [22], and maximum likelihood methods like IQ-TREE [27], PhyML [18], or RAxML [33]. All these methods rely on tree proposals: for a given tree, a new tree similar to the current tree is proposed and is accepted if it fulfils certain conditions. If accepted, the current tree is updated to be this new tree and the procedure repeated. Tree rearrangement operations, which apply local changes to a tree, are commonly used for these tree proposals. Most popular are the Subtree Prune and Regraft (SPR) and Nearest Neighbour Interchange (NNI) tree rearrangements, the latter being a version of SPR restricted to being more local.

An SPR operation (or move) cuts an edge of a phylogenetic tree and reattaches the thereby detached subtree at a different position in the tree. This tree rearrangement (or a version of it) is implemented in many tree inference software packages, including maximum likelihood methods [18, 29, 33], Bayesian inference methods [14, 22, 28] and also recent parsimony-based methods that are able to infer large-scale phylogenies [39]. One reason for the popularity of SPR moves in tree search algorithms is that, unlike very local NNI moves, SPR moves can be used to jump across wider regions of treespace, preventing tree search algorithms from getting stuck in local optima [18, 27]. Similarly, it has been shown for Bayesian inference methods that SPR moves, or modifications thereof, can speed up convergence of Markov chains and improve their mixing when used in combination with other operators [20]. There is however a major obstacle for interpreting how well treespace is traversed when using SPR moves for tree search and tree sampling algorithms: computing the SPR distance, i.e. the minimum number of SPR moves required to transform one tree into another, is 𝒩 𝒫-hard [5, 19]. Despite this 𝒩 𝒫-hardness result, fixed-parameter tractable algorithms for computing SPR distances exist [35], which can be used for analysing modes of posterior distributions [36] or for computing supertrees [35, 37]. Understanding properties of the SPR treespace, which can be viewed as a graph where vertices represent trees that are connected by edges if the trees are connected by an SPR operation, has proven useful for getting a better understanding of phylogenetic inference methods [37].

Analysing the geometry of the classic SPR treespace has lead to numerous results for SPR on rooted and unrooted trees [37, 38]. Although SPR operations are defined in the same way for both types of trees and the two resulting treespaces share most of their properties, the techniques to prove those properties differ significantly. 𝒩 𝒫-hardness of the problem of computing distances, for example, has first been shown for the rooted SPR [5] and only three years later for the unrooted case [19]. An important mathematical structure for both of these proofs is the Maximum Agreement Forest (MAF). A MAF for two trees results from deleting a minimum number of edges in each tree so that the two resulting graphs (forests) are isomorphic. The number of connected components of a MAF for two rooted trees coincides with their rooted SPR distance [5]. For unrooted trees, however, this relationship between the SPR distance and the number of connected components of the MAF does not hold [2, 37], which was the reason some erroneous proofs of 𝒩 𝒫-hardness made it to the literature. It also has been found that the maximum distance under both rooted and unrooted SPR is linear in the number of leaves [4, 12], while the number of trees in the 1-neighbourhood, i.e. the number of trees resulting from one SPR move, is quadratic in the number of leaves [2, 31]. The latter is an important property that has been used to determine the curvature of the treespace by Whidden and Matsen [38] in order to analyse mixing properties of Markov Chain Monte Carlo methods, which are used for Bayesian tree inference.

The majority of these results have been derived for trees that do not contain timing information of evolutionary events. We refer to trees where branch lengths represent times, meaning that the evolutionary events represented by internal nodes are dated, as time trees. These time trees are of interest in many applications. Software packages like BEAST [14] and BEAST2 [7], for example, infer time trees and rely on tree proposals that incorporate timing information of evolutionary events. A version of SPR moves for time trees has been introduced by Höhna, Defoin-Platel, and Drummond [20]. The authors analysed the suitability of this move as a tree proposal operator for Bayesian inference using MCMC algorithms and showed it performed better than some of the previous operators in isolation but it was still better to use a combination. As of May 2023, this move is default in BEAST^1^. A guided version of this proposal was introduced in [21], where guiding the choice of destination improved the acceptance ratio. Another version of SPR moves, which works simultaneously on a species tree and multiple gene trees, can be found in the Stacey package for BEAST2 [25]. Even though SPR moves are widely used as tree proposals and have been adapted to work for time trees, little research has gone into SPR treespaces that take times of evolutionary events into account.

A version of SPR moves for time trees that has been studied mathematically is the one introduced by Song [32], where ranked trees are considered. Ranked trees (which are called ordered trees in [32]) are rooted phylogenetic trees with internal nodes ordered according to times of corresponding evolutionary events and leaves are assumed to be sampled at the same time (ultrametric). The SPR moves defined by Song [32] can move a subtree to a different place in the tree under the condition that the rank of the reattachment node is greater than the rank of the root of the moved subtree. Bounds for neighbourhood sizes and diameters are provided [32], but no further results are known. Despite these efforts to investigate SPR moves for ranked trees, there is still a gap for analysing tree inference methods using SPR for time trees, as the version of SPR moves as defined by Song [32] is currently not used in phylogenetic inference.

In this paper we fill this gap by considering an alternative definition of SPR moves for ranked trees, where we require the height (i.e. rank) at which a subtree is cut and reattached to be the same. This restriction on the rank reattachment is inspired by ranked SPR moves in phylogenetic inference software: The Fixed Node Prune and Regraft move introduced in Höhna, Defoin-Platel, and Drummond [20] as tree proposal for Bayesian inference methods is the extension of our ranked SPR moves to time trees. We call these tree rearrangements Horizontal SPR (HSPR) moves and the corresponding treespace HSPR. Motivated by the importance of this move in computational phylogenetics, we study mathematical properties of this treespace focusing on the metric space of ranked trees that is given by the HSPR move. Studying this treespace helps us understand its fundamental properties, which is important to analyse tree inference algorithms using HSPR for tree proposals as well as posterior distributions of time trees output by BEAST or BEAST2, similar to how it has been done in [36]. By additionally allowing rank moves, which swap the order of two nodes in a ranked tree, we define a further metric space called RSPR (Ranked SPR). We include rank moves in the same way as it has been done in a variation of NNI introduced in [16] for ranked trees (RNNI) that allows distances to be computed in polynomial time [10], unlike in the classical NNI treespace, where this problem is 𝒩 𝒫-hard. This suggests that the complexity of computing distances can be different in classical SPR and its ranked version. This paper is structured as follows. We introduce notations and define the new treespaces in Section 2 before discussing some of its fundamental properties in Section 3. This especially includes the cluster property, which a treespace possesses if shortest paths between trees preserve shared information in the form of common clusters. This property has proven to be important for the polynomial time algorithm in RNNI [10] and for fixed-parameter tractable algorithms in SPR [26, 37]. The discussion of the cluster property is followed by some observations on the shape of shortest paths and the relationship of RSPR and HSPR shortest paths (Section 4). Finally, we establish a surprising result on how the distance between two trees can change after adding a new leaf to them (Section 5). We provide an open source implementation of horizontal SPR moves [8], which we use to show some properties of ranked SPR spaces computationally.

## 2 Preliminaries

A *rooted binary phylogenetic tree* is a pair *T* = (*t, ϕ*) where *t* is a rooted binary tree and *ϕ* : *L*(*t*) → *X* is a bijective map from the set of leaves *L*(*t*) of *t* to a set of labels *X* = {*l*_1_, *l*_2_, *…, l*_*n*_}. Throughout this paper, we refer to rooted binary phylogenetic tree simply as *rooted trees*, and we assume that all trees have *n* leaves, unless stated otherwise.

Let *T* be a rooted tree that contains an edge *e* = (*u, v*) and let *T* |_*v*_ be the subtree of *T* rooted in *v*. A *subtree prune and regraft* (SPR) move on *e* in *T* transforms this tree into a new rooted binary rooted phylogenetic tree by the following three steps:

i. *Prune the subtree T* |_*v*_: delete the edge *e* from *T*, resulting in two connected components *T* |_*v*_ and *T*_*ρ*_, where *T*_*ρ*_ contains the root *ρ* of *T*.
ii. Suppress the resulting node of degree two in *T*_*ρ*_ (node *u*), so that *T*_*ρ*_ is a rooted tree.
iii. *Reattach T*|_*v*_: either by introducing a new node *w* on an edge *f* in *T*_*ρ*_ and adding an edge (*w, v*), or by adding a new root *ρ*′ and adding edges (*ρ*′, *ρ*) and (*ρ*′, *v*).

### Theorem 1.

*The decision problem* SPR:

*Instance: Two rooted trees T and R and an integer k*

*Question: Is d*_SPR_(*T, R*) ≤ *k?*

*is* 𝒩 𝒫*-complete*.

A proof for this theorem can be found in Bordewich and Semple [5]. This proof relies on the equivalence of the SPR distance between two trees and the size of a maximum agreement forest. A *maximum agreement forest* (MAF) of two rooted trees *T* and *R* can be interpreted as a forest that results from cutting the minimum number of edges from both *T* and *R* to result in the same forest, and its size |MAF(*T, R*)| is defined as the number of its connected components. For a formal definition, see [5]. The equality |MAF(*T, R*)| = *d*_SPR_(*T, R*) implies that the same edge is only pruned once on a shortest SPR path. Furthermore, distance computation can be broken down into smaller problems if two trees share some information in the form of common clusters [26], which we will formally define later.

### 2.1 Ranked Trees

A *ranked binary phylogenetic tree* is a pair *T* = (*T*_*u*_, rank) consisting of a rooted tree *T*_*u*_ and a function *rank* : *V* → {0, 1, *…, n* − 1}, where *V* is the set of nodes of *T*_*u*_, such that: (i) rank(*v*) = 0 if and only if *v* is a leaf, (ii) rank(*v*) ≠ *rank*(*w*) for all internal nodes *v* ≠*w*, and (iii) rank(*v*) *<* rank(*w*) if *w* is on the path from the root of *T*_*u*_ to *v*. We refer to rank(*v*) as the *rank of v*, and we denote node of rank *i* in a ranked binary phylogenetic tree *T* by (*T*)_*i*_. For simplicity of notation, we say *tree* or *ranked tree* to refer to ranked binary phylogenetic trees, unless stated otherwise. The assumption that all leaves have rank 0 can be interpreted as requiring ranked trees to be ultrametric.

If *T* = (*T*_*u*_, rank) is a tree, we call the rooted tree *T*_*u*_ the *unranked version* of *T*, as it can be interpreted as *T* without ranks, but with leaf labels. Two trees *T* and *R* are *identical* if there is a graph isomorphism between them that preserves leaf labels and ranks. We then write *T ≃R*, and if the trees are not identical we write *T* ≄ *R*. An example of a tree *T* with annotated ranks and its unranked version *T*_*u*_ is given in Figure 2.

**Figure 1.**
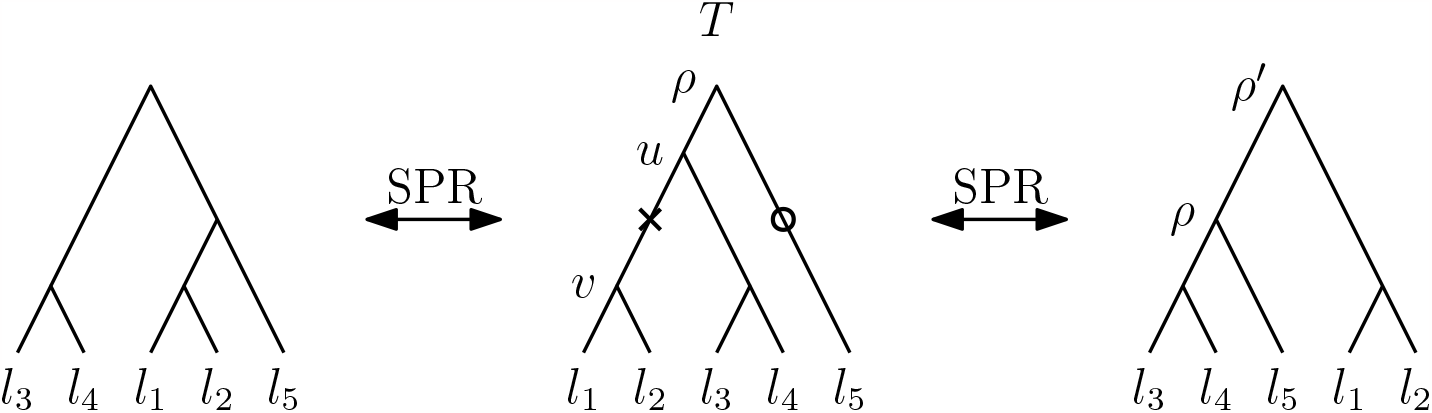
Two rooted SPR moves. The cross marks the edge that is cut to prune the subtree *T* |_*v*_ of *T* with leaf set {*l*_1_, *l*_2_}. In the tree on the left this subtree is reattached on the edge highlighted by a circle, connecting *l*_5_ with its parent, and in the tree on the right the pruned subtree is reattached as child of a newly introduced root *ρ*′.

**Figure 2.**
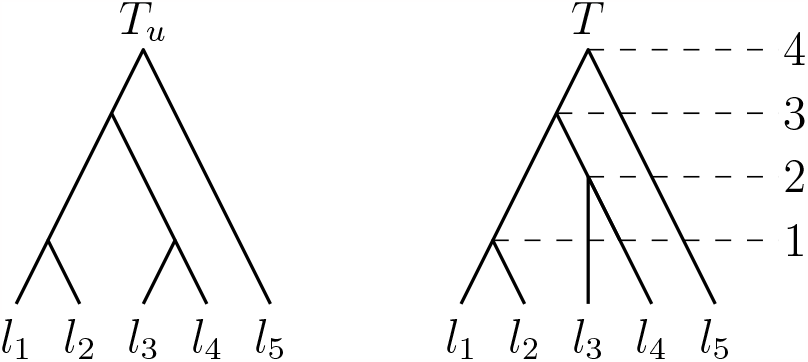
A rooted tree *T*_*u*_ on the left and a ranked tree *T* = (*T*_*u*_, rank) on the right.

A *subtree T* |_*v*_ of a tree *T* is a tree rooted in a node *v* of *T* that contains exactly the nodes that descend from *v* in *T*, annotated by the same ranks as in *T*. We then say that *T* |_*v*_ is *induced* by the node *v*. Note that by this definition of subtrees, the subtree of a ranked tree is not necessarily a ranked tree itself. We denote the parent of a node *v* by parent(*v*) and if a subtree *T* |_*v*_ is induced by *v*, we call parent(*v*) *parent of the subtree T* |_*v*_. To emphasise that we are considering the parent of a subtree *T* |_*v*_ in a tree *T* we might write parent_*T*_ (*T* |_*v*_).

Since we consider trees where internal nodes are assigned unique ranks and leaves are assigned unique leaf labels, we can uniquely identify nodes in a tree by their labels, using the label function

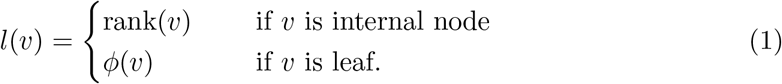

Throughout this paper we assume {1, …, *n* − 1} ∩ {*l*_1_, *…, l*_*n*_} = ∅, which results in *l* being a bijective function. We can therefore uniquely identify a node *v* by its label *l*(*v*), so we will refer to a node *v* simply by *l*(*v*). For example, we might refer to the internal node with rank *i* as *node i* and to the leaf with label *l*_*j*_ as *node l*_*j*_ or *leaf l*_*j*_. We define a strictly partial order ≺_*T*_ on the co-domain of *l* so that *l*(*u*) ≺_*T*_ *l*(*v*) if rank_*T*_ (*u*) *<* rank_*T*_ (*v*). If it is clear that we consider the tree *T* we might simply write ≺ for ≺_*T*_.

Using the label function (1) allows us to represent edges (*u, v*) in a tree as (*l*(*u*), *l*(*v*)). We call the set *E*(*T*) = {(*l*(*v*), (*l*(*w*)) | (*v, w*) is an edge in the tree *T*)} the *edge set of T*. *E*(*T*) uniquely defines *T*. We say that an edge (*l*^+^, *l*^−^) *covers* a rank *i* ∈ {1, *…, n* − 1} if *l*^−^ ≺ *k* ≺ *l*^+^.

A *cluster C* of a tree *T* is the set of leaves descending from an internal node *v* in *T*. We then say that the node *v induces* the cluster *C*. A *cherry* is a subtree consisting of one internal node with two leaves as children, and we refer to a cherry by its cluster containing these two leaves, e.g. {*c*_1_, *c*_2_}. If the internal node of such a cherry has rank *i*, we say that the cherry {*c*_1_, *c*_2_} has rank *i*. Given a set of leaves *S* and a tree *T*, the subtree *T* |_*S*_ of *T induced by S* is the subtree of *T* with minimum number of leaves that contains all leaves of *S*. If *S* is a cluster of *T, T* |_*S*_ contains exactly the leaves of *S*.

We can uniquely represent a tree *T* by a list of its clusters sorted according to increasing rank. This representation is called *cluster representation* and has been introduced by Collienne and Gavryushkin [10]. The leftmost tree in Figure 3 for example has cluster representation [{*l*_1_, *l*_2_}, {*l*_3_, *l*_4_}, {*l*_1_, *l*_2_, *l*_5_}, {*l*_1_, *l*_2_, *l*_3_, *l*_4_, *l*_5_}]. We include the cluster induced by the root in the cluster representation of a tree, even though this cluster is simply the set of all leaf labels {*l*_1_, *l*_2_, *…, l*_*n*_} for all trees on *n* leaves.

**Figure 3.**
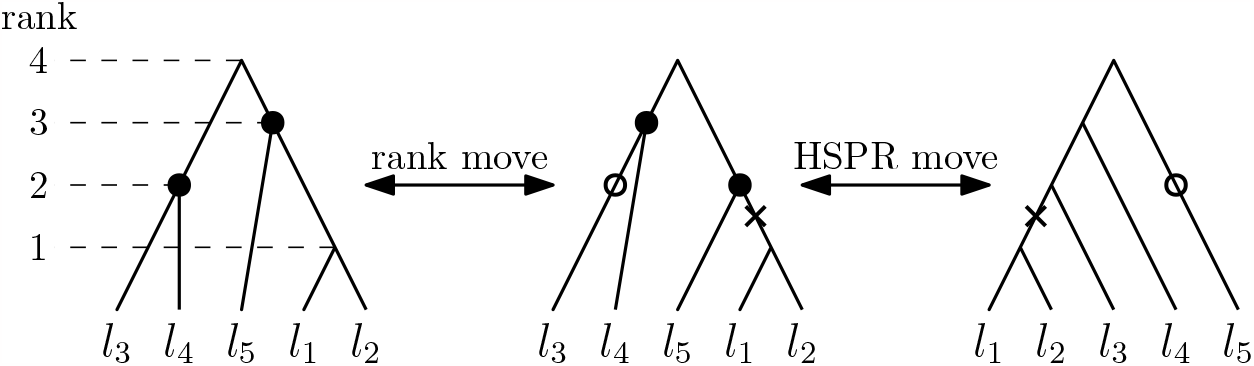
Rank move on the left, swapping the ranks of the nodes highlighted the leftmost tree. On the right an HSPR move at rank 2 is illustrated, moving the subtree induced by {*l*_1_, *l*_2_} by cutting the edge highlighted by a cross and re-attaching it at the edge highlighted by a circle.

### 2.2 Subtree Prune and Regraft for Ranked Trees

A *horizontal* SPR *move* (HSPR move) on an edge *e* of a tree *T*, which cannot be incident to the root, transforms this tree into a tree *T* ′ by shifting the top node of *e* horizontally to another branch. Formally, the move is performed in the following three steps:

i. *prune the subtree T* |_*v*_: delete the edge *e* to obtain two subtrees *T* |_*v*_ and *T*_*ρ*_, where *T*_*ρ*_ has the same root *ρ* as *T*,
ii. suppress the resulting node of degree 2 in *T*_*ρ*_ (top node of edge *e*), whose rank we denote by *i*,
iii. *reattach T* |_*v*_ *on an edge f* : reattach the root of *T* |_*v*_ to a newly introduced node *v* at rank *i* on an edge *f* that covers rank *r* in *T*_*ρ*_.

Since the changes done to the tree *T* move node *i*, we call this an HSPR *move at rank i*. A *rank move* on a tree *T* can be applied to two nodes with consecutive ranks if they are not connected by an edge, and swaps the ranks of these two nodes. An example of an HSPR move and a rank move can be found in Figure 3.

Note that HSPR moves and rank moves are reversible and in contrast to SPR moves for rooted trees, HSPR moves do not allow subtrees to be reattached above the root of a tree.

We are now ready to introduce our main objects of study in this paper, two treespaces extending the classical SPR treespace to ranked trees.

The Ranked SPR (RSPR) space is a graph where vertices represent trees on *n* leaves that are connected by an edge if one tree can be transformed into the other by an HSPR move or a rank move. The Horizontal SPR (HSPR) space is a graph where vertices represent trees on *n* leaves that are connected by an edge if one can be transformed into the other by an HSPR move.

We refer to the moves allowed in RSPR space, i.e. rank moves and HSPR moves, as RSPR *moves*.

A *path* between two trees in HSPR (RSPR) is a sequence of trees *p* = [*T*_0_, *T*_1_, *…, T*_*d*_] such that *T*_*i*_ and *T*_*i*+1_ are connected by an HSPR (RSPR) move for all *i*. If a path *p* contains *d* + 1 trees, we say it has *length* |*p*| = *d*. A *shortest path* between trees *T* and *R* is a path of minimal length connecting *T* and *R*. The length of such a path is called the *distance* between trees *T* and *R*, and we refer to this distance as *d*_HSPR_(*T, R*) in HSPR space and *d*_RSPR_(*T, R*) in RSPR space.

### HSPR moves on edge sets

Let us consider how an HSPR move changes the set of edges of a tree. By the definition of HSPR moves, there are four edges involved in an HSPR move at rank *i* on a tree *T* : The three edges incident to the node of rank *i*, and the edge *f* on which the pruned subtree gets reattached. All other edges stay the same in the tree *T* and its HSPR neighbour *T* ′. Let *e* = (*i, l*) be the edge that is cut by the HSPR move between *T* and *T* ′, let (*i*^+^, *i*) and (*i, i*^−^) be the other two edges incident to *i*, and let *f* = (*j*^+^, *j*^−^) be the reattachment edge in *T*. Then the HSPR move between *T* and *T* ′ changes these edges as follows:

i. *prune the subtree T* |_*v*_: delete the edge *e* = (*i, l*)
ii. suppress the resulting node of degree 2: replace edges (*i*^+^, *i*) and (*i, i*^−^) by an edge(*i*^+^, *i*^−^).
iii. *reattach T* |_*v*_: replace edge *f* = (*j*^+^, *j*^−^) by (*j*^+^, *i*) and (*i, j*^−^) and add edge (*i, l*).

An illustration of the trees *T* and *T* ′ with these node labels is provided in Figure 4. The difference between the edge sets of *T* and *T* ′ can be summarised to

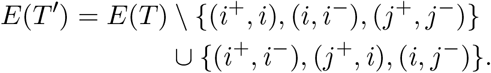

**Figure 4.**
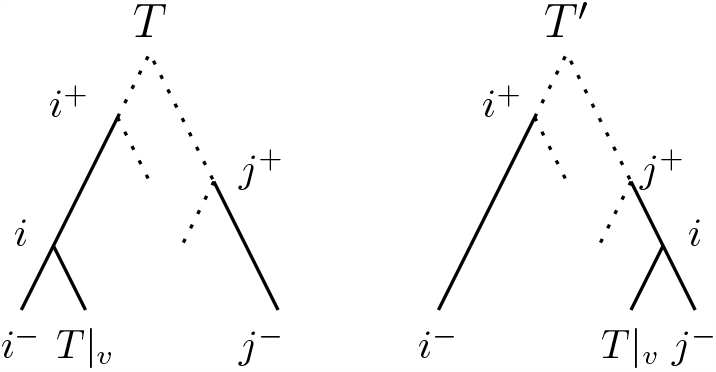
HSPR move pruning the subtree *T* |_*v*_, moving it to the edge (*i*^+^, *i*^*™*^). Dotted edges represent parts of the tree that potentially contain further nodes.

Conversely, if an edge set *E*(*T*′) can be described in this way, then *T* and *T* ′ are connected by an HSPR move:

#### Theorem 2

*Let E*(*T*) *be the set of edges of a tree T*. *A tree T* ′ *with*

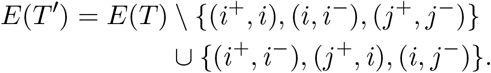

*is* HSPR *neighbour of T if and only if* (*i*^+^, *i*), (*i, i*^−^), (*j*^+^, *j*^−^) ∈ *E*(*T*) *and j*^−^ ≺ *i* ≺ *j*^+^.

Note that *j*^−^ ≺ *i* ≺ *j*^+^ is required in Theorem 2, as the reattachment edge needs to cover the rank *i* of the HSPR move.

Using Theorem 2, we can define an HSPR move between trees *T* and *T* ′ by providing the difference in edge sets *E*(*T*) and *E*(*T*′). We say that a change in edge sets describes a *valid* HSPR move, if the edges that are removed from *E*(*T*) fulfil the conditions of Theorem 2.

### HSPR **moves on cluster representation**

Similar to using edge sets to describe HSPR moves, we can also use the cluster representation to describe an HSPR move between two trees. Let *T* and *R* be trees connected by an HSPR move at rank *i* as described in Theorem 2 and let the clusters induced by *i*^−^, *T* |_*v*_ (the subtree that is moved), and *j*^−^ be *A, B*, and *C*, respectively. Any of *A, B*, and *C* could be just a leaf, e.g. *A* = {*l*_*k*_} for some *k*, but for simplicity we refer to those sets as clusters, too. Let furthermore *T* = [*C*_1_, *C*_2_, *…, C*_*i*−1_, *A* ∪ *B, C*_*i*+1_, *…, C*_*n*−1_] be the cluster representation of *T*. The HSPR move described in Theorem 2 then creates a new cluster *B* ∪ *C* at rank *i*, as in *T* ′ the node *i* has *T* |_*v*_ and *j*^−^ as children. All clusters induced by nodes with rank less than *i* remain unchanged between *T* and *T*′. Since the subtree induced by *B* becomes sibling of the subtree induced by *C* in *T* ′, the move between *T* and *T* ′ removes *B* from every cluster of *T* that contains *B* but not *C* and is induced by a node with rank greater than *i*. On the other side, *B* is added to every cluster induced by a node with rank greater than *i* that contains *C* in *T*. All remaining clusters induced by nodes with rank greater than *i* that do not contain *B* or *C* remain unchanged between *T* and *T* ′.

We can summarise this to describe *T* ′ by its cluster representation

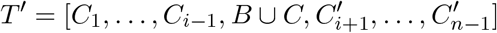

with

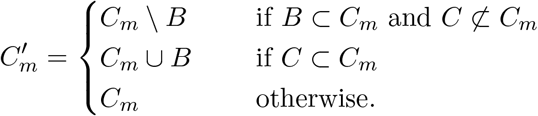

Conversely, if the difference between the cluster representation of *T* and *T* ′ can be described in this way, *T* and *T* ′ are connected by an HSPR move:

#### Theorem 3

*Let T* = [*C*_1_, *C*_2_, *…, C*_*i*−1_, *A* ∪ *B, C*_*i*+1_, *…, C*_*n*−1_] *be a tree and C* = *C*_*l*_ *for some l* ∈ {1, *…, i* − 1}. *A tree*

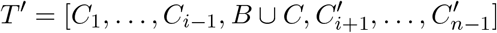

*is connected to T by an* HSPR *move if and only if*

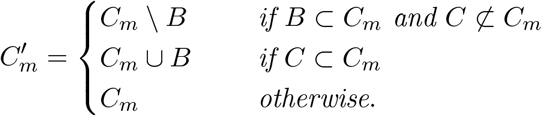

## 3 Basic Properties of HSPR and RSPR

The first question we want to answer is whether our newly defined treespaces are connected (Theorem 4), that is, whether a tree can be transformed into any other tree by a sequence of moves in HSPR or RSPR. Connectedness is essential for these treespaces, as tree rearrangements are used for tree proposals in (MCMC) random walks, which should be able to reach any tree in treespace from any starting tree. For developing and interpreting such random walks it is furthermore important to know how many trees have distance one from a given tree, as well as the maximum distance between any two trees. We establish neighbourhood size (Theorem 5) and maximum distance (diameter) in RSPR and HSPR in Section 3.1. We then investigate the cluster property for RSPR and HSPR and explain the significance of neither of the two spaces having this property (Section 3.2).

### Theorem 4

*The treespaces* HSPR *and* RSPR *are connected*.

*Proof*. We show that any pair of trees *T* ′ and *R* are connected by a path of only HSPR moves. As all HSPR moves are RSPR moves, it then follows that both spaces are connected.

We construct an HSPR path from *T* ′ to *R* by the following bottom up approach, iterating through ranks *k* = 1, *…, n* − 1 of *R*. In every iteration *k*, we perform HSPR moves so that all nodes with ranks less than or equal to *k* induce the same clusters in the tree after iteration *k* and *R*. Let *T* be the tree before iteration *k*, let *R*|_*i*_ and *R*|_*j*_ be the subtrees that are children of the node of rank *k* in *R*, and let *T* |_*l*_ and *T* |_*m*_ be the subtrees that are children of the node of rank *k* in *T*, i.e. parent_*R*_(*R*|_*i*_) = parent_*R*_(*R*|_*j*_) = *k* and parent_*T*_ (*T* |_*l*_) = parent_*T*_ (*T* |_*m*_) = *k*. We can assume that *R*|_*i*_ and *R*|_*j*_ are subtrees in *T*, because we use a bottom up approach that results in all nodes of rank less than *k* inducing the same cluster in *R* and the tree *T* before iteration *k*. Note that in the first iteration *k* = 1, *R*|_*i*_ and *R*|_*j*_ will contain a single leaf only. Consider the following path transforming *T* into a tree *T*_2_ with 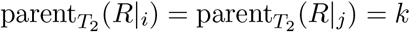 (see Figure 5):

**Figure 5.**
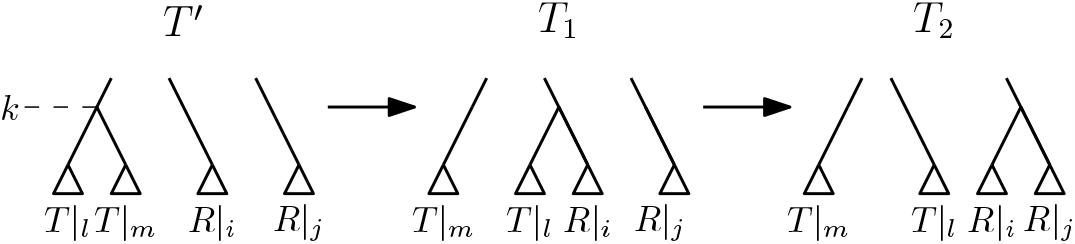
Path between *T* and *T*_2_ as described in the proof of Theorem 4. Only subtrees involved in the moves given in the theorem are shown in this illustration.

First, perform an HSPR move at rank *k* that prunes *T* |_*l*_ from *T* and reattaches it on the edge between *R*|_*i*_ and parent_*T*_ (*R*|_*i*_). In the resulting tree *T*_1_, 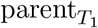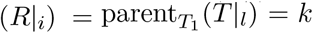. In a second step, we perform an HSPR move pruning the subtree *R*|_*i*_ from *T*_1_ and reattaching it on the edge between *R*|_*j*_ and 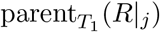, resulting in a tree *T*_2_ with 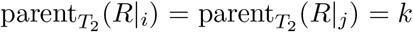. Since all moves between *T* and *T*_2_ are HSPR moves at rank *k*, no clusters induced by nodes less than *k* change between *T* and *T*_2_, while the cluster induced by the node of rank *k* changes so that it coincides between *T*_2_ and *R*. Hence, all cluster induced by nodes with rank less than or equal to *k* are identical in *T*_2_ and *R*. Note that if {*R*|_*i*_, *R*|_*j*_} and {*T* |_*l*_, *T* |_*m*_} intersect, the trees *T*_1_ or *T*_2_ might be equal to *T*, and fewer than two steps are required to reach *T*_2_. At the end of iteration *k*, we update *T* := *T*_2_ and continue with the next iteration *k* + 1. After iteration *n* − 2, we reach *R*, as after each iteration *k* all nodes with rank less than or equal to *k* induce the same clusters in *T* and *R*.

This procedure can be applied to any two trees to compute a path connecting them by a sequence of HSPR moves, which proves the theorem.

The algorithm used to prove Theorem 4 produces a path between any two trees in HSPR, and can hence be used to approximate HSPR distances. An implementation can be found on GitHub [8].

Since HSPR and RSPR are connected undirected graphs, we obtain the following corollary.

### Corollary 1

*d*_HSPR_ *and d*_RSPR_ *are metrics*.

The *(1-)neighbourhood* of a tree *T* in a treespace with distance measure *d* is defined as *NH*(*T*) := {*T* ′ | *d*(*T, T* ′) = 1} and a tree *T* ′ ∈ *NH*(*T*) is called *neighbour* of *T*. We use *NH*_HSPR_(*T*) and *NH*_RSPR_(*T*) to refer to the neighbourhood of *T* in HSPR and RSPR, respectively. Because tree inference algorithms often require sampling tree neighbourhoods, it is important to know the number of *neighbours* of a tree *T* under a tree rearrangement, i.e. |*NH*(*T*)|. In rooted and unrooted SPR the number of neighbours of a tree is quadratic in the number of leaves *n* [2, 31].

*For counting the number of neighbours of a tree in RSPR, we need the following notion: If a tree T* has two nodes *r* and *r* + 1 with rank difference one that are not connected by an edge, we say that [*r, r* + 1] is a *rank interval*. The leftmost tree *T* in Figure 3 for example has two rank intervals: [3, 2] and [2, 1]. We now show that the number of neighbours in RSPR and HSPR is quadratic in the number of leaves, with the number of neighbours in RSPR naturally depending on the shape of the tree. We derive both of these numbers explicitly.

### Theorem 5

*The number of neighbours of a tree T with k rank intervals is* |*NH*_RSPR_(*T*)| = (*n* − 1)(*n* − 2) + *k in* RSPR *and* |*NH*_HSPR_(*T*)| = (*n* − 1)(*n* − 2) *in* HSPR.

*Proof*. The number of neighbouring trees resulting from a rank move on a tree with *k* rank intervals is *k*, as there is one unique rank move for every such interval.

We now count the number of HSPR moves at rank *i* for *i* ∈ {1, *…, n* − 1}. Since every node has two children, two different subtrees can be pruned by an HSPR move at rank *i*. There are *n* − 1 − *i* edges that cover rank *i*, excluding the one on which the node of rank *i* is placed in *T*, so there are *n* − 1 − *i* potential reattachment edges for the pruned subtree. This gives 2(*n* − 1 − *i*) RSPR neighbours of *T* resulting from an HSPR move at rank *i*. And because HSPR moves can be performed at any rank between 1 and *n* − 1, the number of RSPR moves possible on *T* is:

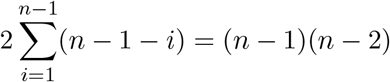

Since all rank moves and HSPR moves result in different trees, *T* has (*n* − 1)(*n* − 2) neighbours in HSPR and (*n* − 1)(*n* − 2) + *k* neighbours in RSPR.

### 3.1 Diameter

The maximum distance between any two trees in a treespace with distance measure *d*, 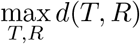, is called the *diameter* of the treespace. When measuring the similarity of trees using a distance metric, knowing the maximum possible distance is essential for interpreting distances. The diameter of both unrooted and rooted SPR space is 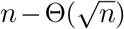 [4, 12] and hence linear in *n*. We show in this section that the diameters of HSPR space and RSPR space are linear in the number of leaves *n*, too, by establishing lower and upper bounds.

#### Corollary 2

*The diameters of* HSPR *space and* RSPR *space have an upper bound of* 2(*n* − 2).

*Proof*. We can use the algorithm introduced in the proof of Theorem 4 to compute a path between any two trees *T* and *R* in HSPR. Since every HSPR move is an RSPR move, the length of this path is an upper bound of the diameter of both HSPR and RSPR space. The algorithm uses a bottom-up approach that, starting at tree *T*, constructs in iteration *k* = 1, *…, n* − 2 the cluster induced by the node of rank *k* in *R*, using at most two HSPR moves. After iteration *n* − 2, we receive the destination tree *R* after at most 2(*n* − 2) HSPR moves. Because the path *p* constructed by this algorithm has length at most 2(*n* − 2), this provides an upper bound to the diameter of HSPR and RSPR.

It is important to note that the algorithm described in the proof of Theorem 4 approximates HSPR distances and does not compute the exact distance for all pairs of trees, which we can show using our implementations [8].

We can also prove a lower bound for the distance between any two trees in HSPR, but first we need the following lemma.

#### Lemma 1

*Let T and R be trees containing x leaves whose parents have different ranks in T and R, i*.*e*. |{(*u, v*) | *v is leaf and* (*u, v*) ∈ *E*(*T*) *and* (*u, v*) ∈ *E*(*R*)}| = *x. Then* 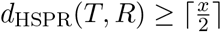.

*Proof*. As described in the technical introduction, an HSPR move at rank *i* between trees *T* and *T* ′ leads to the following difference in edge sets for some edges (*i*^+^, *i*), (*i, i*^−^), (*j*^+^, *j*^−^) ∉ *E*(*T*) where (*i*^+^, *j*^−^) covers rank *i*:

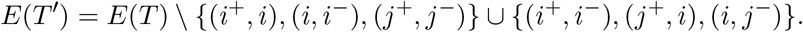

Therefore, the only nodes whose parents change by this HSPR move are the nodes *i, i*^−^, and *j*^−^. Not all of them need to have different a parent after the move, since for example *i*^+^ = *j*^+^ results in the parent of *i* having the same rank in *T* and *T* ′. With (*i*^+^, *i*) and (*i, i*^−^) being edges in *T*, it follows that *i* is an internal node, so only *i*^−^ and *j*^−^ can be leaves. Therefore, an HSPR move can change the parents of at most two leaves.

Since the ranks of *x* parents of leaves differ between *T* and *R* and any HSPR move can fix at most two of those, there are in total at least 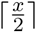 HSPR moves needed to connect *T* and *R*.

The lower bound given in Lemma 1 can be tight. For example, the two leftmost trees in Figure 6 are connected by one HSPR move and the parents of *x* = 2 leaves (*l*_2_ and *l*_5_) have different ranks in the two trees.

**Figure 6.**
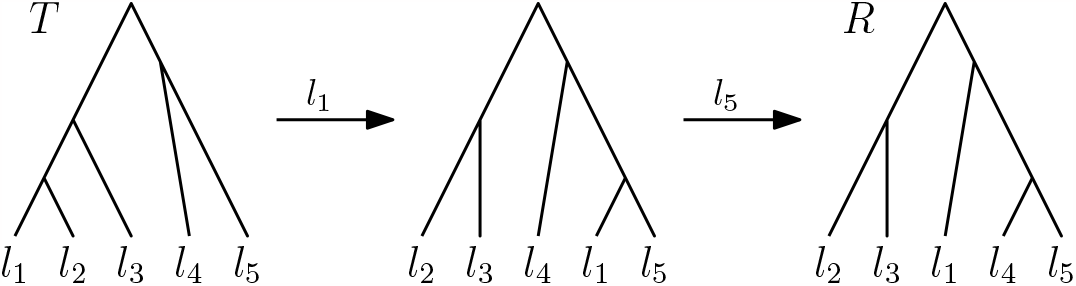
Trees *T* and *R* of the counterexample to the weak cluster property in RSPR and HSPR in Theorem 7. The labels of the arrows indicate leaves that are pruned in the corresponding HSPR moves.

#### Theorem 6

*There are trees T and R with distance* 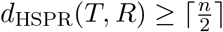.

*Proof*. Let *T* and *R* be the following caterpillar trees:

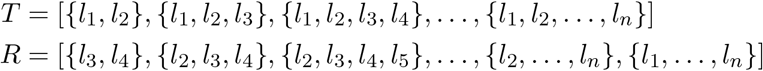

The parent of *l*_1_ and *l*_2_ has rank one in *T*, but not in *R*, where *l*_3_ and *l*_4_ are children of the node of rank one. For all other leaves *l*_*i*_ with *i* ∈ {5, *…, n*}, the rank of the parent also is different in *T* than in *R*: parent_*T*_ (*l*_*i*_) = *i* − 1 *i* − 2 = parent_*R*_(*l*_*i*_). Therefore, the parents of all *n* leaves have different ranks in *T* and *R*, and by Lemma 1 it follows 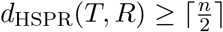.

By Corollary 2 and Theorem 6, the diameter of the HSPR space is linear in *n*. We will see later (Corollary 6) that Theorem 6 also applies to RSPR.

### 3.2 No Cluster property

Two trees *T* and *R share* a cluster *C* if both of them contain *C* as cluster. If on a path *p* every tree contains the cluster *C*, we say that *p preserves* the cluster *C*. We moreover say that a treespace has the *strong cluster property* (simply called *cluster property* by Collienne et al. [9]) if for two trees sharing a cluster *C*, every shortest path between them preserves *C*. In other words, if two trees share some evolutionary information in form of a cluster, this information is preserved along every shortest path between them if the treespace has the cluster property. If for any two trees sharing a cluster *C* there exists a shortest path that preserves *C* in a tree space, we say that this treespace has the *weak cluster property*. Note that the difference between the weak and the strong cluster property is that for the strong cluster property we require all shortest paths to preserve clusters, while for the weak cluster property only one shortest path needs to preserve a shared cluster.

The classic rooted (unranked) version of SPR space has the weak cluster property, which is shown as part of the proof of Theorem 2.2 in [26], where the problem of computing the SPR distance is split into the problem of computing distances for subtrees induced by shared clusters. The weak cluster property of SPR is essential for fixed-parameter tractable algorithms [26, 35] and facilitates proofs for 𝒩 𝒫-hardness of computing SPR distances, as it is related to the formulation of this problem as an agreement forest problem.

Here we show that RSPR and HSPR space have neither the weak nor the strong cluster property. This observation is important as it suggests that the proving technique for 𝒩 𝒫-hardness for rooted (unranked) SPR, which uses maximum agreement forests, cannot be used for ranked trees.

#### Theorem 7

HSPR *space and* RSPR *space do not have the weak cluster property*.

*Proof*. We prove this theorem by considering trees *T* and *R* that share a cluster but have no shortest path between them that preserves this cluster. Let *T* and *R* be the following two trees (see Figure 6):

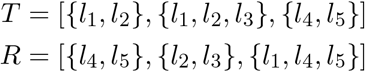

On any path between *T* and *R* preserving the shared cluster {*l*_4_, *l*_5_}, the rank of the node inducing this cluster needs to decrease from 3 in *T* to 1 in *R*. Note that originating from *T*, no HSPR move can decrease the rank of this cluster, the only possible moves preserving {*l*_4_, *l*_5_} are rank moves. There is hence no shortest path between *T* and *R* in HSPR that preserves {*l*_4_, *l*_5_}, i.e. HSPR does not have the weak cluster property.

In RSPR, two rank moves are necessary to decrease the rank of the node inducing {*l*_4_, *l*_5_} to 1, resulting in a tree *R*′ = [{*l*_4_, *l*_5_}, {*l*_1_, *l*_2_}, {*l*_1_, *l*_2_, *l*_3_}]. Since *R*′ is not identical to *R*, further RSPR moves are needed to receive *R*, resulting in a path of length greater than two.

There is however a path from *T* and *R* with only two HSPR moves, where first the leaf *l*_1_ is pruned and moved to the edge (parent(*l*_5_), *l*_5_) and then *l*_5_ is pruned and moved to the edge (parent(*l*_4_), *l*_4_) (see Figure 6). Since this path is shorter than any path in RSPR preserving the cluster {*l*_4_, *l*_5_}, RSPR space does not have the weak cluster property.

## 4 Shortest paths

One of the main hindrances of using tree rearrangement based distance measures is that most of them are 𝒩 𝒫-hard to compute. This applies to rooted and unrooted SPR distance [5, 19] as well as NNI distance [11]. A recent modification of NNI tree rearrangements for ranked trees has however shown to be computationally tractable [10]. Whether the same is true for ranked SPR is a natural question, and we approach it by investigating the structure of shortest paths in RSPR and HSPR treespace. Though the complexity of computing distances in HSPR and RSPR remains unknown, our results in this section are a first step towards understanding shortest paths in these treespaces. In particular, we investigate the relationship of shortest paths in HSPR and RSPR, which hints at the complexity of the shortest path problem in the two treespaces being identical. Gaining insights into the structure of shortest paths might furthermore be useful for developing algorithms for computing or approximating ranked SPR distances.

### 4.1 HSPR Shortest Paths

Shortest paths between trees in HSPR are generally not unique, and we show here that they can be arranged so that the rank at which subtrees are cut does not decrease on a shortest path (Theorem 8). We then use this result to study scenarios at which clusters are preserved along shortest paths: First, we show that if the cherry at rank one is identical in two trees, it is preserved on every shortest path (Theorem 9). Then, by further generalising this result, we show in Corollary 4 that for two trees with identical clusters up to a certain rank, all shortest paths between them preserve these clusters. These results provide insights into the shape of shortest paths in HSPR, and in future research we hope to leverage these results to prove the complexity of computing distances in this treespace.

To prove that there is a shortest path in HSPR on which ranks of HSPR moves do not decrease, we need the following lemma.

#### Lemma 2

*Let p* = [*T, T* ′, *R*] *be a path in* HSPR *such that T and T* ′ *are connected by an* HSPR *move at rank k and T* ′ *and R are connected by an* HSPR *move at rank i with k > i. Then there is a path p*′ = [*T, T* ″, *R*] *where an* HSPR *move at rank i connects T and T* ″ *and an* HSPR *move at rank k connects T* ″ *and R*.

*Proof*. As described in Theorem 2, we can describe the HSPR move between trees *T* and

*T* ′ by the change in the set of edges *E*(*T*), compared to *E*(*T*′):

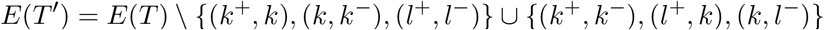

where (*k*^+^, *k*), (*k, k*^−^), (*l*^+^, *l*^−^)∈ *E*(*T*) and (*l*^+^, *l*^−^) covers rank *k*. Similarly, we assume that *E*(*T*′) and *E*(*R*) are related in the following way:

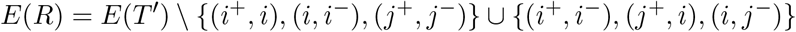

where (*i*^+^, *i*), (*i, i*^−^), (*j*^+^, *j*^−^) ∈ *E*(*T*′) and (*j*^+^, *j*^−^) covers rank *i*. In total, we get

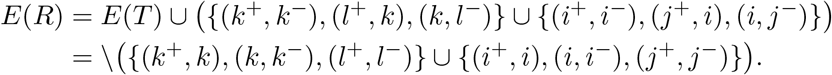

Note that there might be edges that are added to *T* ′ and then deleted in *R*. Therefore, when summarising multiple HSPR moves as set operations, we assume that we consider multisets, even though the edge set of a tree does not contain an edge multiple times. Let

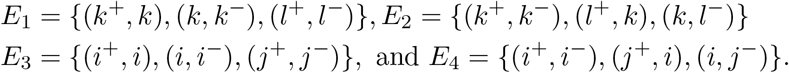

Then *E*_3_ ∩ *E*_1_ = ∅, as otherwise the HSPR move between *T* ′ and *R* would not be possible.

If *E*_2_ ∩ *E*_3_ = ∅, then *E*_3_ ⊂ *E*(*T*), and we can create a tree *T* ″ with *E*(*T* ″) = *E*(*T*) \ *E*_3_ ∪ *E*_4_, i.e. *T* and *T* ″ are connected by an HSPR move at rank *i*. With *E*_2_ ∩ *E*_3_ = ∅ we get *E*_2_ ∈ *E*(*T* ″) and we can perform an HSPR move at rank *k* on *T* ″ that results in *E*(*R*^*\**^) = *E*(*T* ″) \ *E*_2_ ∪ *E*_1_.

Hence, *E*(*R*^*\**^) = *E*(*T*)∪(*E*_3_ ∪*E*_1_)\(*E*_4_ ∪*E*_2_) = *E*(*R*). Therefore, *R*^*\**^ and *R* are identical and the path *p ′*= [*T, T*″, *R*] contains an HSPR move at rank *i* followed by an HSPR move at rank *k*.

We now distinguish different cases in which *E*_2_ ∩ *E*_3_ ∅. With *i*^−^, *j*^−^ ≺ *i* ≺ *k* ≺ *k*^+^, it is (*i, i*^−^) ∉ *E*_2_ and (*l*^+^, *k*) ∉ *E*_3_. Therefore, *E*_2_ ∩ *E*_3_ ≠ ∅ if and only if {(*i*^+^, *i*), (*j*^+^, *j*^−^)} ∩ {(*k*^+^, *k*^−^), (*k, l*^−^)} ∅. We distinguish six different cases, based on this intersection. In every case, we construct an alternative path *p*′ = [*T, T* ″, *R*], where *T* and *T* ″ are connected by an HSPR move at rank *I* and *T* ″ and *R* are connected by an HSPR move at rank *k*. To show that the moves that we describe by edge set changes are valid HSPR moves, we need to show that the edge set changes we provide fulfil the criteria listed in Theorem 2. By our assumption of the moves on *p*, we know that *E*_1_ ⊂ *E*(*T*) and if *e* ∈ *E*_3_ \ *E*_2_, then *e* ∈ *E*(*T*).

1 (*i*^+^, *i*) = (*k*^+^, *k*^−^) and (*j*^+^, *j*^−^) ≠ (*k, l*^−^): We perform an HSPR move on *T* ′ to receive the tree *T* ″ with

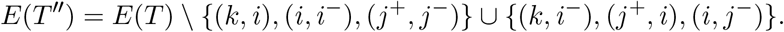

This is a valid move because:

- (*k, i*) ∈ *E*(*T*), because (*k, i*) = (*k, k*^−^) and (*k, k*^−^) ∈ *E*_1_.
- (*i, i*^−^) ∈ *E*(*T*), because (*i, i*^−^) ∈ *E*_3_ and (*i, i*^−^) ∉ *E*_2_.
- (*j*^+^, *j*^−^) ∈ *E*(*T*), because (*j*^+^, *j*^−^) ∈ *E*_3_, and (*j*^+^, *j*^−^) ∉ *E*_2_
- (*j*^+^, *j*^−^) covers rank *i* by the definition of the moves on *p*.

We can then receive a tree *R*^*\**^ by an HSPR move on *T* ″ with

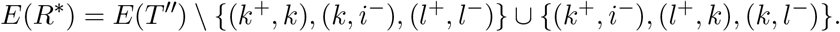

This is a valid move because:

- (*k*^+^, *k*) ∈*E*(*T* ″), because (*k*^+^, *k*) ∈ *E*_1_ and (*k*^+^, *k*) has not been removed between *T* and *T* ″.
- (*k, i*^−^) ∈ *E*(*T* ″), as it has been added between *T* and *T* ″.
- (*l*^+^, *l*^−^)∈ *E*(*T* ″), because (*l*^+^, *l*^−^) ∈ *E*_1_, and (*l*^+^, *l*^−^) has not been removed between *T* and *T* ″.
- (*l*^+^, *l*^−^) covers rank *k* by the definition of the moves on *p*.

With *i*^+^ = *k*^+^ and *i* = *k*^−^ it follows *E*(*R*^*\**^) = *E*(*R*), and hence *R*^*\**^ ≃ *R*.

2 (*i*^+^, *i*) = (*k*^+^, *k*^−^) and (*j*^+^, *j*^−^) = (*k, l*^−^): Let *T* ″ be resulting from *T* by an HSPR move at rank *i* that changes *E*(*T*) as follows:

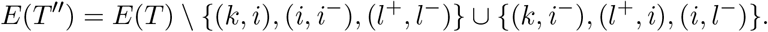

This is a valid move because:

- (*k, i*) ∈ *E*(*T*), because (*k, i*) = (*k, k*^−^) and (*k, k*^−^) ∈ *E*_1_.
- (*i, i*^−^) ∈ *E*(*T*), because (*i, i*^−^) = (*k*^−^, *i*^−^) ∈ *E*_3_ and (*k*^−^, *i*^−^) ∉ *E*_2_.
- (*l*^+^, *l*^−^) ∈ *E*(*T*), because (*l*^+^, *l*^−^) ∈ *E*_1_.
- (*j*^+^, *j*^−^) covers rank *i* by the definition of the moves on *p*.

We can then receive a tree *R*^*\**^ by an HSPR move on *T* ″ with

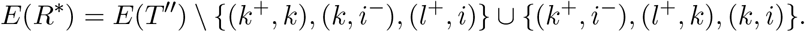

This is a valid move because:

- (*k*^+^, *k*) ∈*E*(*T* ″), because (*k*^+^, *k*) ∈*E*_1_ and (*k*^+^, *k*) has not been removed between *T* and *T* ″.
- (*k, i*^−^) ∈ *E*(*T* ″), as it has been added between *T* and *T* ″.
- (*l*^+^, *i*) ∈ *E*(*T* ″), as it has been added between *T* and *T* ″.
- (*l*^+^, *i*) covers rank *k* by the definition of the moves on *p*.

With *i*^+^ = *k*^+^, *i* = *k*^−^, *j*^+^ = *k*, and *j*^−^ = *l*^−^ it follows *E*(*R*^*\**^) = *E*(*R*) and hence *R*^*\**^ ≃ *R*.

3. (*i*^+^, *i*) = (*k, l*^−^) and (*j*^+^, *j*^−^)≠ (*k*^+^, *k*^−^):

We can perform an HSPR move at rank *i* on *T* that gives us the following tree *T* ″:

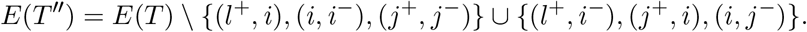

This is a valid move because:

- (*l*^+^, *i*) ∈ *E*(*T*), because (*l*^+^, *i*) = (*l*^+^, *l*^−^) ∈ *E*_1_.
- (*i, i*^−^) ∈ *E*(*T*), because (*i, i*^−^) ∈ *E*_3_ and (*i, i*^−^) ∉ *E*_2_.
- (*j*^+^, *j*^−^) ∈ *E*(*T*), because (*j*^+^, *j*^−^) ∈ *E*_3_ and (*j*^+^, *j*^−^) ∉ *E*_2_.
- (*j*^+^, *j*^−^) covers rank *i* by the definition of the moves on *p*.

We can then perform an HSPR move at rank *k* on *T* ″ to get a tree *R*^*\**^ with

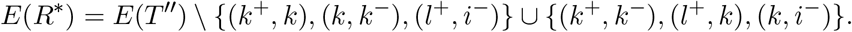

This is a valid move because:

- (*k*^+^, *k*) ∈ *E*(*T* ″), because (*k*^+^, *k*) ∈ *E*_1_ and (*k*^+^, *k*) has not been removed between *T* and *T* ″.
- (*k, k*^−^)∈ *E*(*T* ″), because (*k*^+^, *k*) ∈ *E*_1_ and (*k*^+^, *k*) has not been removed between *T* and *T* ″.
- (*l*^+^, *i*^−^) ∈ *E*(*T* ″), as it has been added between *T* and *T* ″.
- (*j*^+^, *j*^−^) covers rank *k* by the definition of the moves on *p*.

By *i*^+^ = *k* and *i* = *l*^−^ it then follows *E*(*R*^*\**^) = *E*(*R*) and hence *R*^*\**^ ≃ *R*.

4. (*i*^+^, *i*) = (*k, l*^−^) and (*j*^+^, *j*^−^) = (*k*^+^, *k*^−^):

We can perform an HSPR move at rank *i* on *T* to get a tree *T* ″ with

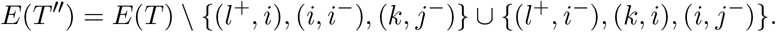

This is a valid move because:

- (*l*^+^, *i*) ∈ *E*(*T*), because (*l*^+^, *i*) = (*l*^+^, *l*^−^) ∈ *E*_1_.
- (*i, i*^−^) ∈ *E*(*T*), because (*i, i*^−^) ∈ *E*_3_ and (*i, i*^−^) ∉ *E*_2_.
- (*k, j*^−^) ∈ *E*(*T*), because (*k, j*^−^) = (*k, k*^−^) ∈ *E*_1_.
- (*k, j*^−^) covers rank *i* by the definition of the moves on *p*.

An HSPR move at rank *i* can transform *T* ″ to a tree *R*^*\**^ with

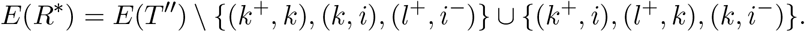

This is a valid move because:

- (*k*^+^, *k*) ∈*E*(*T* ″), because (*k*^+^, *k*) ∈ *E*_1_ and (*k*^+^, *k*) has not been removed between *T* and *T* ″.
- (*k, i*) ∈ *E*(*T* ″), as it has been added between *T* and *T* ″.
- (*l*^+^, *i*^−^) ∈ *E*(*T* ″), as it has been added between *T* and *T* ″.
- (*l*^+^, *i*^−^) covers rank *k* by the definition of the moves on *p*.

By *i*^+^ = *k, i* = *l*^−^, *j*^+^ = *k*^+^, and *j*^−^ = *k*^−^ it follows *E*(*R*^*\**^) = *E*(*R*) and therefore *R*^*\**^ ≃ *R*.

5. (*j*^+^, *j*^−^) = (*k*^+^, *k*^−^) and (*i*^+^, *i*)≠ (*k, l*^−^):

Performing an HSPR move at rank *i* on *T* can give us a tree *T* ″ with

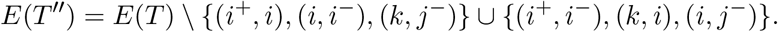

This is a valid move because:

- (*i*^+^, *i*) ∈ *E*(*T*), because (*i*^+^, *i*) ∈ *E*_3_ and (*i*^+^, *i*) ∉ *E*_2_.
- (*i, i*^−^) ∈ *E*(*T*), because (*i, i*^−^) ∈ *E*_3_ and (*i, i*^−^) ∉ *E*_2_.
- (*k, j*^−^) ∈ *E*(*T*), because (*k, j*^−^) = (*k, k*^−^) ∈ *E*_1_.
- (*k, j*^−^) covers rank *i* by the definition of the moves on *p*.

An HSPR move at rank *k* can therefore convert *T* ″ into a tree *R*^*\**^ with

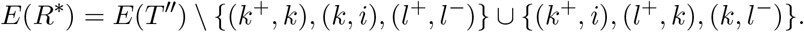

This is a valid move because:

- (*k*^+^, *k*) ∈*E*(*T* ″), because (*k*^+^, *k*) ∈*E*_1_ and (*k*^+^, *k*) has not been removed between *T* and *T* ″.
- (*k, i*) ∈ *E*(*T* ″), as it has been added between *T* and *T* ″.
- (*l*^+^, *l*^−^) ∈*E*(*T* ″), because (*l*^+^, *l*^−^) ∈*E*_1_ and (*l*^+^, *l*^−^) has not been removed from *T* to get *T* ″.
- (*l*^+^, *l*^−^) covers rank *k* by the definition of the moves on *p*.

By *j*^+^ = *k*^+^ and *j*^−^ = *k*^−^ it follows *E*(*R*^*\**^) = *E*(*R*) and hence *R*^*\**^ ≃ *R*.

6. (*j*^+^, *j*^−^) = (*k, l*^−^) and (*i*^+^, *i*)≠ (*k*^+^, *k*^−^):

We can perform an HSPR move at rank *i* on *T* to get a tree *T* ″ with edge set

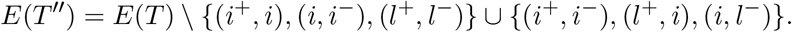

This is a valid move because:

- (*i*^+^, *i*) ∈ *E*(*T*), because (*i*^+^, *i*) ∈ *E*_3_ and (*i*^+^, *i*) ∉ *E*_2_.
- (*i, i*^−^) ∈ *E*(*T*), because (*i, i*^−^) ∈ *E*_3_ and (*i, i*^−^) ∉ *E*_2_.
- (*l*^+^, *l*^−^) ∈ *E*(*T*), because (*l*^+^, *l*^−^) ∈ *E*_1_.
- (*l*^+^, *l*^−^) covers rank *i* because *i* ≺ *k* ≺ *l*^+^ and *j*^−^ = *l*^−^ ≺ *i*.

Then an HSPR move at rank *k* is possible on *T* ″ and transforms this tree into *R*^*\**^ with

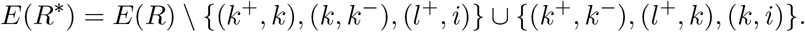

This is a valid move because:

- (*k*^+^, *k*) ∈*E*(*T* ″), because (*k*^+^, *k*) ∈*E*_1_ and (*k*^+^, *k*) has not been removed between *T* and *T* ″.
- (*k, k*^−^)∈*E*(*T* ″), as (*k, k*^−^) ∈*E*_1_ and (*k, k*^−^) has not been removed between *T* and *T* ″.
- (*l*^+^, *i*) ∈ *E*(*T* ″), because it has been added by the move between *T* and *T* ″.
- (*l*^+^, *i*) covers rank *k* by the definition of the moves on *p*.

By *j*^+^ = *k* and *j*^−^ = *l*^−^, it follows *E*(*R*\*) = *E*(*R*), hence *R*^*\**^ ≃ *R*.

#### Theorem 8

*There is always a shortest* HSPR *path between any two trees T and R on which the rank of* HSPR *moves increases monotonically*.

*Proof*. Let *p* be a shortest path between trees *T* and *R*. By Lemma 2 we can take any pair of consecutive HSPR moves at ranks *k* and *i* and if *k > i*, we can replace them by two HSPR moves so that first a rank *i* and then a rank *k* HSPR move is performed. By replacing all such pairs of HSPR moves iteratively, we receive a path on which the ranks of HSPR moves increase monotonically.

To show that all shortest paths preserve a shared cherry at rank one (Theorem 9), we need the following two lemmas, which describe HSPR moves at a fixed rank *k* along a shortest path. These lemmas are interesting on their own as they are informative about the local geometry of the HSPR treespace. Recall that (*T*)_*i*_ is the node of rank *i* in *T* whereas *T* |_*i*_ is the subtree induced by that node.

#### Lemma 3

*Let T and R be trees containing subtrees T* |_1_, *T* |_2_, *…, T* |_*d*_ *with T* |_*j*_≄ *T* |_*m*_ *for all j* ≠ *m, so that* 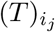*and* 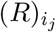 *induce the subtree T* |_*j*_ *for j* = 1, *…, d in T and R,respectively. Let* 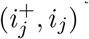 *be the edges between* 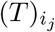 *and its parent in T for all j* = 1, *…, d. If*

i. 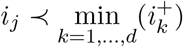*for all j* = 1, *…, d and*
ii. 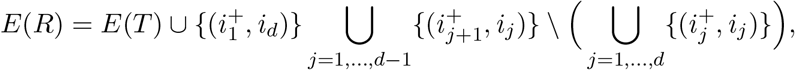

*Then d*_HSPR_(*T, R*) ≤ *d*.

Informally, the difference between *T* and *R* is the positioning of the subtrees

*T* |_1_, *T* |_2_, *…, T* |_*d*_, which we can interpret as a permutation of these subtrees. By changing every edge 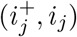 of *T* to 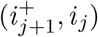 in *R*, every subtree *T* |_*j*_ in *R* is attached where *T* |_*j*+1_ is in *T* for all *j* = 1, *…, d* − 1, and with 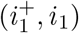 being replaced by 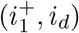, *T* |_*d*_ in *R* is attached where *T* |_1_ is in *T*. We could describe this permutation of the subtrees by (*T* |_1_, *T* |_2_, *…, T* |_*d*_), using the cycle notation for permutations.

*Proof*. We assume without loss of generality that 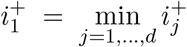. If otherwise 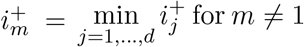, we rename the subtrees *T* |_((*j*+*d*−*m*+1) mod *d*)+1_ to *T* |_*j*_ for all *j* = 1, *…, d*.

By the assumptions of the lemma, all edges 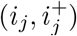 with *j*≠ *i* in *T* cover rank 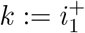. Let 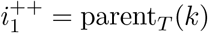 and let *i*_0_ be the sibling of *i*_1_ in *T*, i.e. parent_*T*_ (*i*_0_) = parent_*T*_ (*i*_1_) = *k*.

We now create a path *p* = [*T*_0_ ≃ *T, T*_1_, *…, T*_*d*_ ≃ *R*] of length *d*. We describe the moves on *p* iteratively for every pair *T*_*j*−1_, *T*_*j*_ by using the edge set notation. For every move we make, we prove that it is a valid move by showing that the conditions of Theorem 2 are fulfilled. Note that we can infer from *T* |_*j*_≄ *T* |_*m*_ that *i*_*j*_≠ *i*_*m*_ for all *j m*.

We define the first move on *p* by the following change of the edge set of *T* :

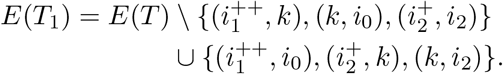

This is a valid move, because:

- 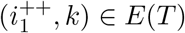 by definition of 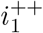 and *k*.
- (*k, i*_0_) ∈ *E*(*T*) by definition of *k* and *i*_0_.
- 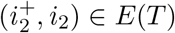 by the assumption of the lemma.
- 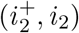 covers rank *k* by the assumptions of the lemma.

The next moves on *p* between *T*_*j*−1_ and *T*_*j*_ for *j* = 2, *…, d* − 1 are described by the following changes in edge sets:

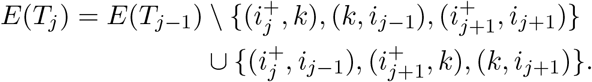

These are valid moves, because:

- 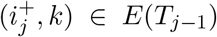, because this edge is added to *E*(*T*_*j*−1_) by the previous move between *T*_*j*−2_ and *T*_*j*−1_.
- (*k, i*_*j*−1_) ∈ *E*(*T*_*j*−1_), because this edge is added in the move from *T*_*j*−3_ to *T*_*j*−2_, and it is not changed when moving from *T*_*j*−2_ to *T*_*j*−1_.
- 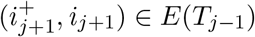, because 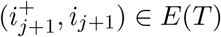 by the assumptions of the lemma, and with *i*_*j*_≠ *i*_*m*_ for all *j*≠ *m* and 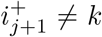 for all *j*≠ 1, this edge is not changed on any previous move on *p*.
- 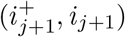covers rank *k* by the assumption of the lemma.

The last step, transforming *T*_*d*−1_ to *T*_*d*_, we apply an HSPR move to *T*_*d*−1_ that changes the edge set as follows:

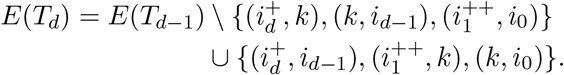

This is a valid move, because:

- 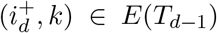, because this edge is added to *E*(*T*_*d*−1_) by the previous move between *T*_*d*−2_ and *T*_*d*−1_.
- (*k, i*_*d*−1_) ∈ *E*(*T*_*d*−1_), because this edge is added in the move from *T*_*d*−3_ to *T*_*d*−2_, and it is not changed when moving from *T*_*d*−2_ to *T*_*d*−1_.
- 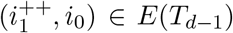, because it is added to *E*(*T*_1_) in the first move on *p* and with *i*_*j*_≠ *i*_*m*_ for all *j*≠ *m* and 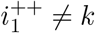, we can infer that it is not removed from the edge set of any tree on any previous move on *p*.
- 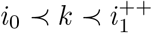 by definition of *k*.

We can summarise all changes to the edge sets along *p* using multisets, which can be simplified using 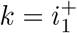:

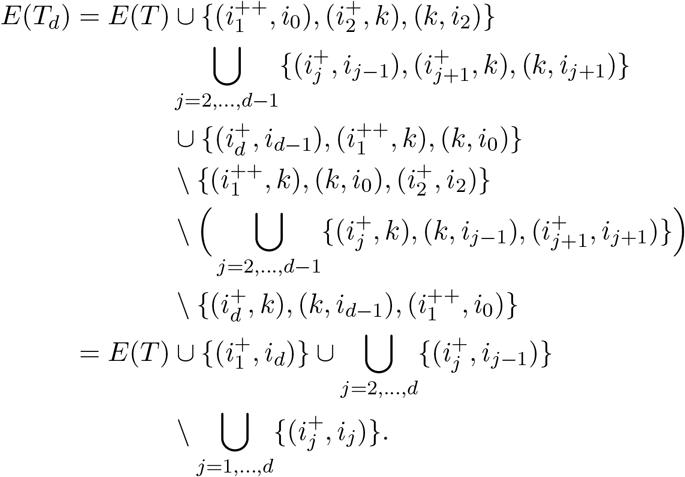

It follows *E*(*T*_*d*_) = *E*(*R*) and therefore, *T*_*d*_ ≃ *R*. Hence, *p* is a path from *T* to *R* of length *d*, proving *d*_HSPR_(*T, R*) ≤ *d*.

Another interesting property of shortest paths is that of Lemma 4: No subtree can move twice by HSPR moves at the same rank. It is however possible for one subtree to move twice at different ranks on a shortest path. For example in Figure 7, the subtree only consisting of the leaf *a*_5_ is moved by the first HSPR move at rank one and by the last HSPR move at rank two.

**Figure 7.**
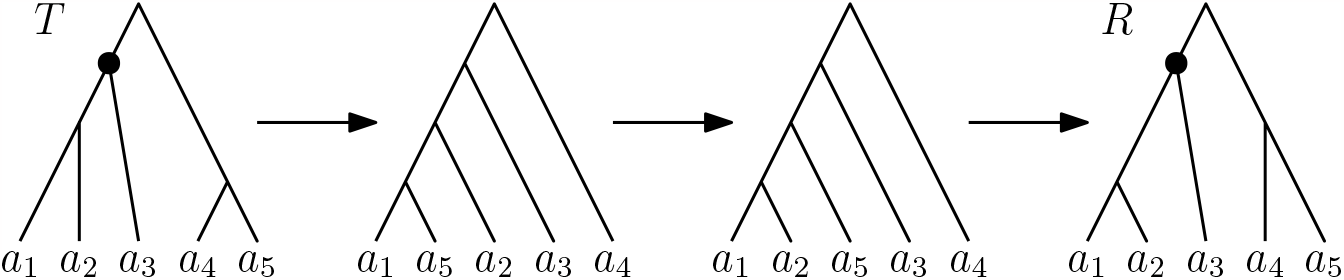
Shortest HSPR path between trees *T* and *R*. The subtree consisting of only the leaf *a*_5_ is moved by the first and the last HSPR move. The path displayed here does not preserve the shared cluster {*a*_1_, *a*_2_, *a*_3_} at rank three. The nodes inducing this cluster are highlighted in *T* and *R*. On this path only subtrees consisting of single leaves move.

#### Lemma 4

*Let T and R be trees connected by a shortest path p containing only* HSPR *moves at rank k, moving subtrees* 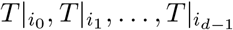 *in this order, i*.*e. p has length d*.

*Then every* 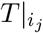 *is subtree of both T and R and* 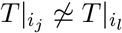*for all j* ≠ *l* ∈ {0, 1, *…, d* − 1}.

*Proof*. Let *p* be a shortest path only containing HSPR moves at rank *k* as described in the lemma. Every subtree 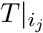 with *j* ∈ {0, 1, *…, d* − 1} is induced by a node *i*_*j*_ with rank less than *k* in *T*. Therefore, no edges inside any of these subtrees can change on *p*, which implies that 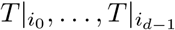 are subtrees in all trees on *p*, including *T* and *R*.

To show 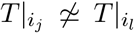for all *j* ≠ *l*, we assume to the contrary that a subtree 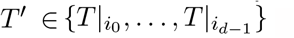 moves twice on a *p*. To simplify notation, we assume without loss of generality that it is 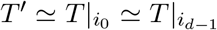 and that no other subtree moves twice on *p*. Note that it could be *d* − 1 = 1, in which case *T* ′ moves twice in two consecutive moves.

Let 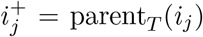 in *T*. Since the first move on *p* moves the subtree 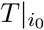 and all moves on *p* are HSPR moves at rank *k*, it is *i*_0_ = *k*. Furthermore, the subtree 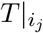 needs to have node *k* as its parent before it gets pruned from a tree on *p*, as all moves are HSPR moves at rank *k*. This implied that 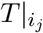 gets reattached on the edge 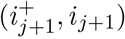 for every *j* = 0, *…, d* − 2 to ensure that the next move can prune the subtree 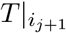at rank *k*. Let *i*_0_ be the sibling of *i*_1_ in *T*, i.e. parent_*T*_ (*i*_0_) = parent_*T*_ (*i*_1_) = *k* and let 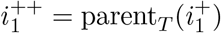.

Assuming *p* = [*T*_0_ ≃ *T, T*_1_, *…, T*_*d*_ ≃ *R*], the HSPR moves along *p* can formally be described by the following changes in edge sets (see Theorem 3):

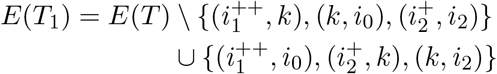

and for *j* = 2, *…, d* − 2:

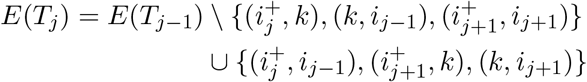

By our assumptions on *p*, 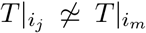 for all *j≠ m* in {1, *…, d* − 1}, we know that 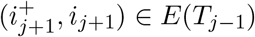 for all *j* = 1, *…, d* − 1. We can use similar arguments to those used in the proof of Lemma 3 to show that the changes of edge sets presented here describe valid HSPR moves up to *T*_*d*−2_.

For the last move on *p*, moving the subtree 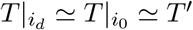 by an HSPR move at rank *k*, the subtree 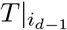 needs to become sibling of 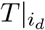 by an HSPR move at rank *k* between *T*_*d*−2_ and *T*_*d*−1_. The edge connecting *i*_1_, which is the root of 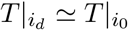, and its parent is 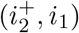, which is created by the move between *T*_1_ and *T*_2_ and with 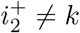 and *i*_*j*_ ≠ *i*_*k*_ for all *j* ≠ *k* in {1, *…, d* − 1}, this edge is not changed again on *p* until reaching *T*_*d*−2_. The move between *T*_*d*−2_ and *T*_*d*−1_ can therefore be described by the following change in the edge set of *T*_*d*−2_:

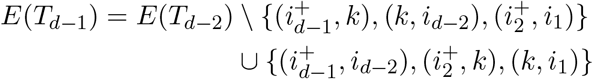

Let 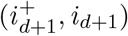 be the edge on which 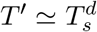gets reattached by the last move on *p*. We can then write this last move as

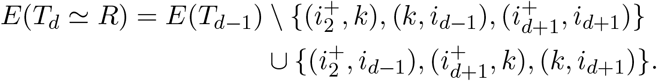

It is again not hard to see with Theorem 2 that this describes a valid HSPR move.

Using multisets, we can describe the difference between *E*(*R*) and *E*(*T*) as follows:

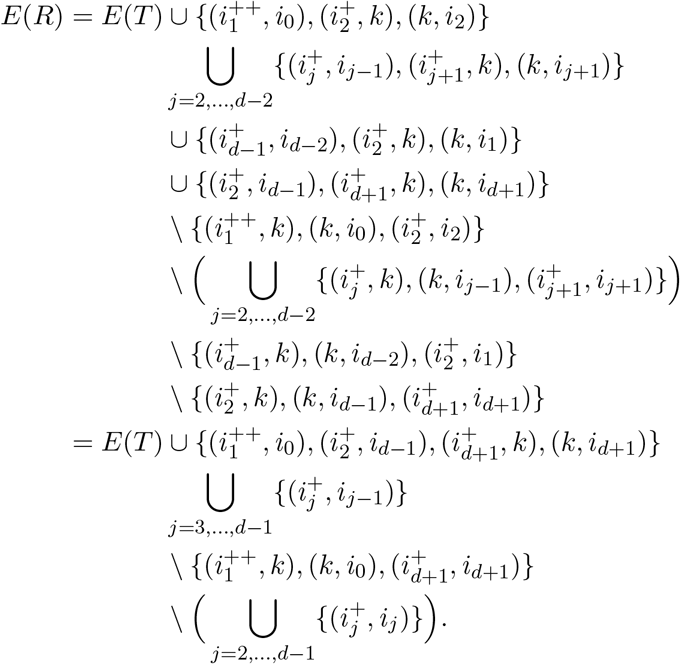

Based on this, we can define an alternative path from *T* to *R*. Therefore, we define a neighbour 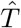of *T* by

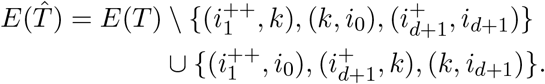

This describes a valid HSPR move on *T*, because:

- 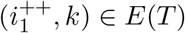 by the definition of 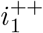 and *k*.
- (*k, i*_0_) ∈ *E*(*T*) by the definition of *k* and *i*_0_.
- 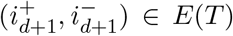, because 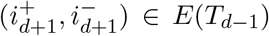 by the definition of moves on *p* and this edge is not deleted from the edge set on any move on *p* before *T*_*d*−1_ is reached.
- 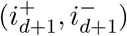 covers rank *k* by the assumption that all HSPR move are HSPR moves at rank *k*.

Knowing the difference between *E*(*T*) and *E*(*R*), we can summarise the difference between *E*(*R*) and 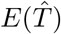 to:

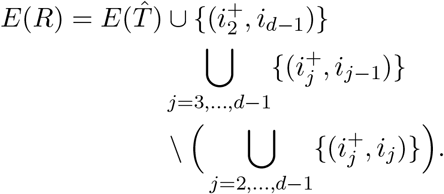

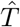 and *R* fulfil the assumptions of Lemma 3, so we get 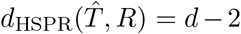. Therefore, the path from *T* to *R* via 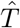 has length 1 + *d* − 2 = *d* − 1 and is shorter than *p*, contradicting our assumption that *p* is a shortest path. Hence, no subtree is moved twice on a shortest with only HSPR moves at fixed rank *k*.

We are now ready to show that a shared cherry at rank one is preserved on every shortest path.

#### Theorem 9

*Let T and R be trees and x, y* ∈ {*l*_1_, *…, l*_*n*_} *so that both T and R have* {*x, y*} *as cherry at rank one. Then every tree on every shortest path between T and R has* {*x, y*} *as cherry at rank one*.

*Proof*. For this proof we assume to the contrary that *T* and *R* are connected by a shortest path *p* containing a tree whose cherry at rank one is not {*x, y*}. We furthermore assume that *T* and *R* have minimum distance among all tree pairs connected by such a shortest path. It then follows that *p* contains HSPR moves at rank one only: Otherwise, we could change the order of ranks of HSPR moves along the path to have all rank one HSPR moves first (Lemma 2). Since the cherry at rank one cannot change by HSPR moves at rank greater than one, *T* and *R* would then not be a minimum distance counterexample. Therefore, all HSPR moves on *p* are at rank one and hence move subtrees that consist of one leaf only.

By the assumption on *T* and *R*, either *x* or *y* is pruned by the first move on *p*. We assume that *x* is moved first, otherwise we swap notations for *x* and *y*. Since the last move on *p* reconstructs the cherry {*x, y*} at rank one, and the subtree containing only the leaf *x* can only move once at rank one (Lemma 4), the last leaf that moves on *p* is *y*. Let *a*_0_ = *x, a*_1_, *a*_2_, *…, a*_*d*−1_, *a*_*d*_ = *y* be the sequence of leaves moved on *p*. By Lemma 4, all these leaves are distinct i.e. *a*_*i*_ ≠ *a*_*j*_ for all *i* ≠ *j*. Note that *d* − 1 ≥ 1, as after the first move on *p* the parent of *y* has rank greater than one, but the last move on *p* moves *y* by an HSPR move at rank one, for which the parent of *y* needs to have rank one. Moving *a*_*d*−1_ ≠ *x* to become sibling of *y* in the second to last move is hence necessary to get to *R*, so *d* − 1 ≥ 1. The path *p* as described above is depicted in Figure 8.

**Figure 8.**
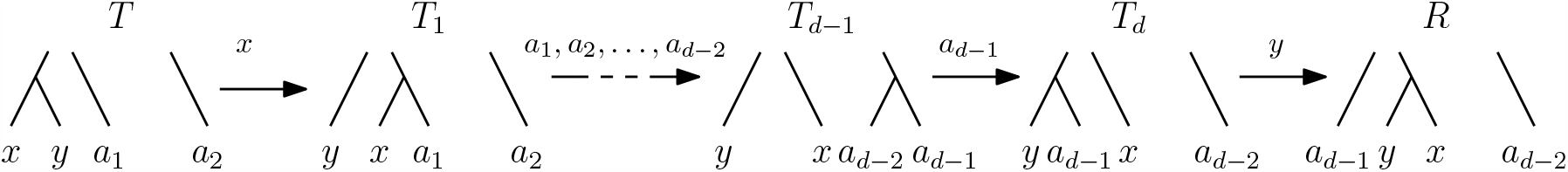
Trees *T* and *R* with cherry {*x, y*} at rank one and all moves on a path *p* from *T* to *R* that does not preserve the cherry as described in the proof of Theorem 9. The labels of the arrows indicate the leaves that are pruned and reattached by the corresponding HSPR moves.

Let 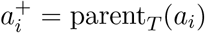 for all *i* = 1, *…, d* and *p*_*xy*_ = parent_*T*_ (1). Note that this implies 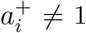 for all *i*. Assuming *p* = [*T*_0_ ≃ *T, T*_1_, *…, T*_*d*_, *T*_*d*+1_ ≃ *R*], we can then describe the moves on *p* similar to how it has been done in the proof of Lemma 4:

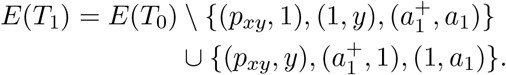

This describes a valid move, since:

- (*p*_*xy*_, 1) ∈ *E*(*T*_0_) by the definition of *p*_*xy*_.
- (1, *y*) ∈ *E*(*T*_0_), because *y* is in the cherry of rank one.
- 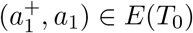 by the definition of 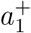.
- 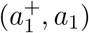 covers rank 1, because *a*_1_ is a leaf and 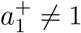.

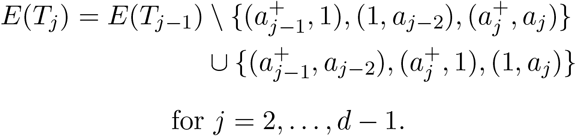

This describes a valid move, because:

- 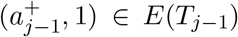, because it has been added to this set by the previous move between *T*_*j*−2_ and *T*_*j*−1_.
- (1, *a*_*j*−2_) ∈ *E*(*T*_*j*−1_), because it has been added to *E*(*T*_*j*−2_) by the move between *T*_*j*−3_ and *T*_*j*−2_ and with *a*_*i*_ ≠ *a*_*j*_ for all *i* ≠ *j*, this edge has not been removed from *E*(*T*_*j*−2_) to get *E*(*T*_*j*−1_).
- 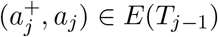, because 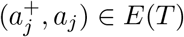and with *a*_*i*_ ≠ *a*_*j*_ for all *i*≠ *j*, this edge is not removed from the edge set in any previous move on *p*.
- 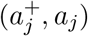 covers rank 1, because *a*_*j*_ is a leaf and 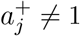.

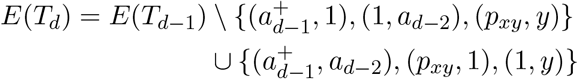

This describes a valid move, because:

- 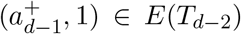, because it has been added to this set by the previous move between *T*_*d*−3_ and *T*_*d*−2_.
- (1, *a*_*d*−2_) ∈ *E*(*T*_*d*−2_), because it has been added to *E*(*T*_*d*−3_) by the move between *T*_*d*−2_ and *T*_*d*−2_.
- (*p*_*xy*_, *y*) ∈ *E*(*T*_*d*−2_), because it has been added to *E*(*T*_1_) by the first move on *p* and with *y* ≠ *a*_*i*_, 1 for all *i*, this edge is not removed from the edge set in any previous move on *p*.
- (*p*_*xy*_, *y*) covers rank 1, because (*p*_*xy*_, 1) and (1, *y*) are edges in *T* .

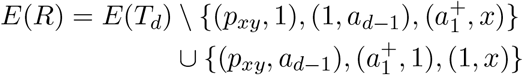

This describes a valid move, because:

- (*p*_*xy*_, 1) ∈ *E*(*T*_*d*_), because it has been added to this set by the previous move between *T*_*d*−1_ and *T*_*d*_.
- (1, *a*_*d*−1_) ∈ *E*(*T*_*d*_), because it has been added to *E*(*T*_*d*−2_) by the move between *T*_*d*−2_ and *T*_*d*−1_ and with *a*_*i*_ ≠ *a*_*j*_ for all *i* ≠ *j*, this edge has not been removed from *E*(*T*_*d*−1_) to get *E*(*T*_*d*_).
- 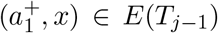, because it has been added to *E*(*T*_2_) by the second move on *p* (*x* = *a*_0_) and with *x* ≠ *a*_*i*_, 1 for all *i*, this edge is not removed from the edge set in any previous move on *p*.
- 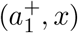 covers rank 1, because *x* is a leaf and 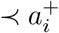 for all *i*.

Since some edges that are added along *p* are later deleted, we can describe the changes between *E*(*T*) and *E*(*R*) by using multisets:

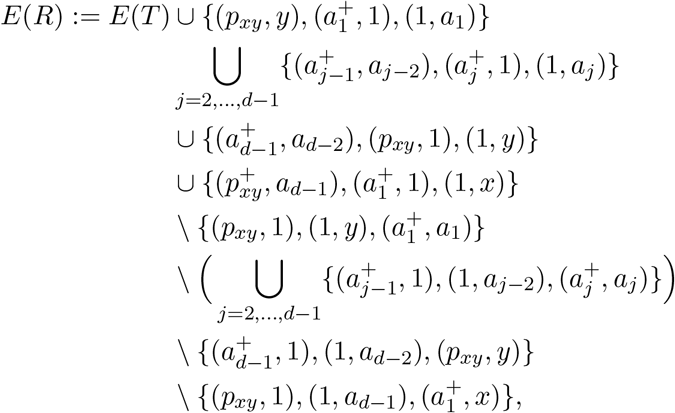

which we can summarise to:

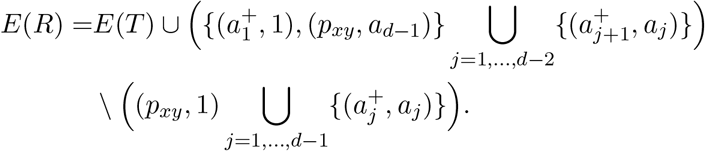

*T* and *R* fulfil the requirements of the trees of Lemma 3, where 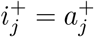 and *i*_*j*_ = *a*_*j*_ for all *j* = 1, *…, d* − 1, 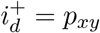, and *i*_*d*_ = 1. Applying this lemma gives us *d*_HSPR_(*T, R*) ≤ *d*.

Therefore, *p* is not shortest paths, contradicting our assumption that there is a shortest path from *T* to *R* that does not preserve the shared cherry.

Theorem 9 implies that the distance of two trees with identical cherry at rank one does not change when one of the leaves in this cherry is deleted (Corollary 3). We will see in Theorem 11 that we can generally not assume that the distance stays the same or decreases when deleting a leaf from two trees.

#### Corollary 3

*Let T and R be trees on n leaves sharing their cherry {c*_1_, *c*_2_} *at rank one and let T* ′ *and R*′ *result from deleting c*_2_ *from T and R, respectively, suppressing the internal node of rank one and subtracting one from the rank of all remaining internal nodes so that T and R are trees on n* − 1 *leaves. Then d*_HSPR_(*T* ′, *R*′) = *d*_HSPR_(*T, R*).

*Proof*. By Theorem 9 all shortest HSPR paths from *T* to *R* preserve the cherry {*c*_1_, *c*_2_}. Therefore, deleting *c*_2_ from every tree on a shortest *T* -*R*-path, suppressing the resulting degree-two nodes, and subtracting one from the ranks of all remaining nodes, gives a path between *T* ′ and *R*′, i.e. *d*_HSPR_(*T* ′, *R*′)≤ *d*_HSPR_(*T, R*). If there was a path between *T* ′ and *R*′ that was shorter than *d*_HSPR_(*T, R*), then adding a leaf *c*_2_ as sibling of *c*_1_ with parent of rank one and adding one to the ranks of all other internal nodes in every tree on *p* results in a path between *T* and *R* of length less than *d*_HSPR_(*T, R*), which is a contradiction. Hence, *d*_HSPR_(*T, R*) = *d*_HSPR_(*T* ′, *R*′).

Another observation that follows from Theorem 9 is that if the lowest part of two trees, i.e. all clusters up to a certain rank, is identical, then no shortest path in HSPR changes this part of the tree.

#### Corollary 4

*Let T and R be trees so that for some i* ∈ {2, *…, n*} *the cluster induced by* (*T*_*′*_)_*j*_ *is identical to the cluster induced by* (*R*)_*j*_ *for all j < i. Then every node* (*T*)_*j*_ *in every tree T* ′ *on every shortest path between T and R induces the same cluster as* (*T*)_*j*_ *and* (*R*)_*j*_ *for all j < i*.

*Proof*. To prove this corollary, we assume to the contrary that there is a node with rank less than *i* that induces the same cluster in *T* and *R*, but this cluster is not present on a shortest path *p* between *T* and *R*. Let *j* be the rank of such a node so that there is no other node with this property in *T* and *R* with rank less than *j*. Then all clusters induced by nodes of rank less than *j* are present in all trees on *p*. We can iteratively apply Corollary 3 to the trees *T* and *R* where in the first iteration deleting a cherry leaf and updating the rest of the tree as described in the lemma results in trees *T*_1_ and *R*_1_ with *d*_HSPR_(*T, R*) = *d*_HSPR_(*T*_1_, *R*_1_). This can be repeated until trees *T*_*j*−1_, *R*_*j*−1_ are received with *d*_HSPR_(*T*_*j*−1_, *R*_*j*−1_) = *d*_HSPR_(*T, R*), where *T*_*j*−1_ and *R*_*j*−1_ are trees on *n*−(*j* −1) leaves. As all clusters induced by nodes of rank less than *j* are present in all trees on *p*, the leaves that are deleted from *T* and *R* to receive *T*_*j*−1_ and *R*_*j*−1_, respectively, can be deleted in the same order in all trees on *p* in the same way, and we receive a path *p*′ from *T*_*j*−1_ to *R*_*j*−1_ with |*p*′| = |*p*|. Since the cluster induced by the nodes of rank *j* is the same in *T* and *R, T*_*j*−1_ and *R*_*j*−1_ have the same cherry at rank one. Because *p* does not preserve the cluster at rank *j*, there must be a tree on *p*′ not containing the cherry of rank one in *T*_*j*−1_ and *R*_*j*−1_, which implies by Corollary 3 that *p*′ is not a shortest path, i.e. *d*_HSPR_(*T*_*j*−1_, *R*_*j*−1_) *<* |*p*′| = |*p*|. This however is a contradiction to *d*_HSPR_(*T*_*j*−1_, *R*_*j*−1_) = *d*_HSPR_(*T, R*) = |*p*|. Therefore, there cannot be a shortest path *p* between *T* and *R* that does not preserve the cluster of rank *j < i* that is present in *T* and *R*, which concludes the proof of this corollary.

It is in general not true that a cluster *C* that is induced by nodes of the same rank *r* in two trees *T* and *R* is present in all trees on all shortest paths. A counterexample to this can be found in Figure 7 where the shared cluster {*a*_1_, *a*_2_, *a*_3_} at rank three is not present in every tree on any shortest path from *T* to *R*, which we found out through exhaustive search using our implementation [8].

### 4.2 RSPR Shortest Paths

The RSPR space can be interpreted as an extension of HSPR in which rank moves are added to provide shortcuts between some trees. Here, we investigate how the addition of rank moves changes shortest paths. Our analysis of these paths will provide insights into the relationship of the complexity of computing distances in HSPR and RSPR. We show in Theorem 10 that we can change the order of moves on paths in RSPR, while not changing their length, so that all rank moves are grouped at the beginning followed by a sequence of only HSPR moves. This indicates that shortest paths in HSPR and RSPR can be very similar, sometimes identical (e.g. in the case of caterpillar trees, Corollary 6), and suggests that the complexity of computing shortest paths (and distances) might be the same in both treespaces. We first need the following lemma:

#### Lemma 5

*Let T and R trees and p* = [*T, T* ′, *R*] *a path with an* HSPR *move between T and T* ′ *and a rank move between T* ′ *and R. Then there is a path p*′ = [*T, T* ″, *R*] *consisting of either two* HSPR *moves or a rank move followed by an* HSPR *move*.

Whether the path *p*′ in Lemma 5 contains two HSPR moves or a rank move followed by an HSPR move depends on the specific moves on *p*, as we will see in the proof of this lemma. For this proof we use the cluster representation of trees and describe HSPR moves by the corresponding changes in cluster representation as described in Theorem 3.

*Proof*. Let *k* and *k* + 1 for some *k* ∈ {1, *…, n* − 3} be the ranks of the nodes of the rank move between *T* ′ and *R*. Let *T* ′ = [*C*_1_, *…, C*_*k*−1_, *A* ∪*B, C* ∪*D, C*_*k*+2_, *…, C*_*n*−1_] be the cluster representation of *T* ′, which is illustrated in the top middle of Figure 9. Since *T* ′ and *R* are connected by a rank move of nodes *k* and *k* + 1, the cluster representation of *R* is *R* = [*C*_1_, *…, C*_*k*−1_, *C* ∪ *D, A* ∪ *B, C*_*k*+2_, *…, C*_*n*−1_] (see top right of Figure 9).

**Figure 9.**
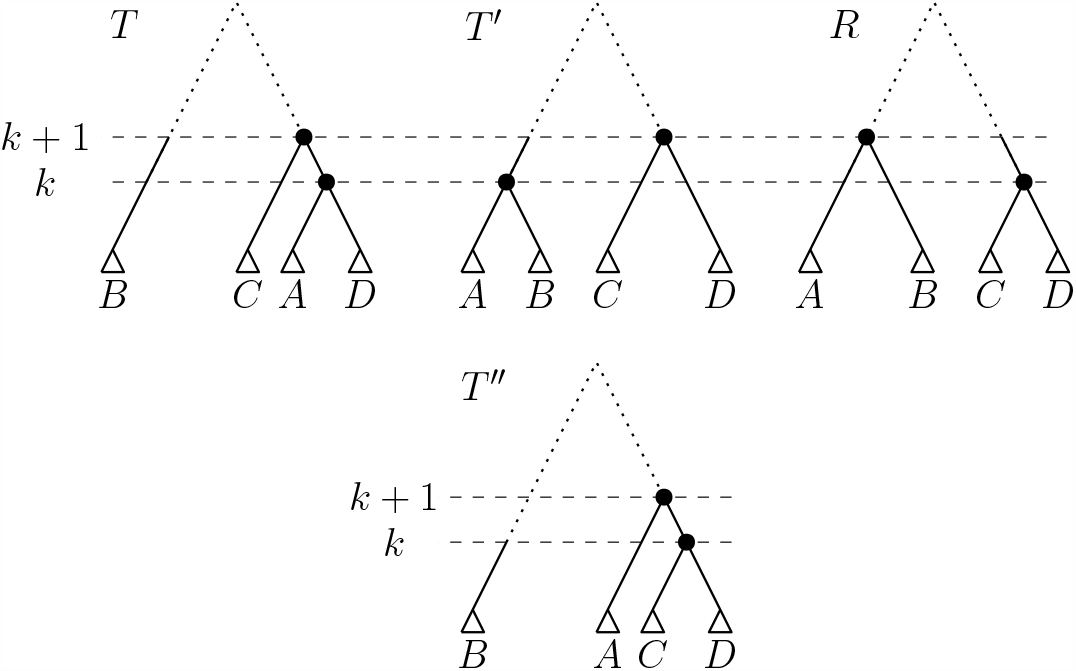
Trees *T, T* ′, and *R* on *p*, and alternative path *p*′ from *T* to *R* via *T* ″ at the bottom if *T* and *T* ′ are connected by an HSPR move at rank *k* and the nodes of rank *k* and *k* + 1 in *T* are connected by an edge, as explained in Figure 2.1. The dotted parts of the trees might contain further nodes and leaves.

In the following, we distinguish different HSPR moves possible between *T* and *T*′ and show how to replace *T* ′ by a tree *T* ″ to get a path *p*′ = [*T, T* ″, *R*] with either two HSPR moves or a rank move followed by an HSPR move.

1. The HSPR move between *T* and *T* ′ is neither a rank *k* nor a rank *k* + 1 HSPR move. An HSPR move at rank greater than *k* does not change the clusters induced by nodes *k* and *k* + 1, so we can in this case first perform a rank move of nodes *k* and *k* + 1 on *T* to get a tree *T* ″. Since all clusters of rank greater than *k* + 1 are identical in *T* and *T* ″ we can perform an HSPR move on *T* ″ that changes the clusters of this tree in exactly the same way as they change between *T* and *T* ′. This HSPR move on *T* ^*′′*^ then results in *R*, giving us a path *p*′ = [*T, T* ″, *R*] with the desired properties. If the HSPR move between *T* and *T* ′ is an HSPR move at rank *r < k*, it might change the clusters induced by the nodes of rank *k* and *k* + 1. It does however not matter, in which order these two clusters appear in the tree, they would change in the exact same way if they were swapped. Therefore, we can first swap the ranks of the nodes with rank *k* and *k* + 1 in *T*, resulting in a tree *T* ″, and then perform the same HSPR move on *T* ″ as the one between *T* and *T* ′. This results in a path *p*″ = [*T, T* ″, *R*] with a rank move between *T* and *T* ″ and an HSPR move between *T* ″ and *R*.
2. *T* and *T* ′ are connected by an HSPR move at rank *k*. Without loss of generality we can assume that *T* |_*A*_ is moved between *T* and *T* ′, otherwise we swap notations for *A* and *B*. We now further distinguish whether there is an edge connecting the nodes of rank *k* and *k* + 1 in *T*.
  2.1 Let there be an edge connecting the nodes with ranks *k* and *k* + 1 in *T*. Note that by our assumptions on *T* ′, parent_*T*′_ (*T* |_*C*_) = parent_*T*′_ (*T* |_*D*_) = *k* + 1 and parent_*T*′_ (*T* |_*A*_) = *k*. Therefore, an HSPR move at rank *k* on *T* ′ that moves the subtree *T* |_*A*_ and creates a tree *T* containing an edge (*k* + 1, *k*) must move *T* |_*A*_ to become sibling of either *T* |_*C*_ or *T* |_*D*_. We assume without loss of generality that *T* |_*A*_ and *T* |_*D*_ are siblings in *T*, as depicted in Figure 9, otherwise we change notation for *C* and *D*. Then the cluster representation of *T* is:

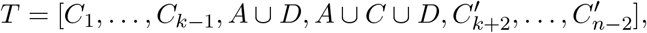

where for all *m* = *k* + 1, *…, n* − 2

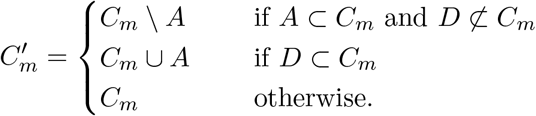

To get a path *p*′ with the desired properties, we perform an HSPR move at rank *k* on *T* that moves *T* |_*D*_ to become sibling of *T* |_*C*_, resulting in a tree *T* ″ in which the parent of *T* ″|_*A*_ has rank *k* + 1. This tree *T* ″ has cluster representation:

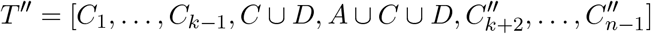

where for all *m* = *k* + 1, *…, n* − 2:

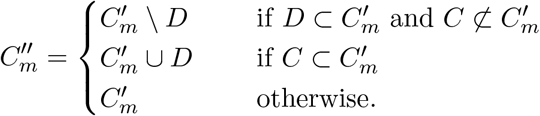

Remember that every cluster is the union of two clusters at lower rank and/or leaves of a tree. Since the node (*T*)_*k*+1_ induced the cluster *A* ∪ *C* ∪ *D*, it is 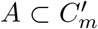 if and only if 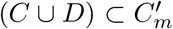 for all *m* ≥ *k* + 1. Therefore, all clusters induced by nodes with rank greater than *k* + 1 are the same in *T* and *T* ″: 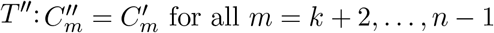. We can now perform an HSPR move at rank *k* + 1 on *T* ″ that moves the subtree *T* ″|_*A*_ to become sibling of *T* ″|_*B*_, which results in a tree 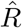 with cluster representation:

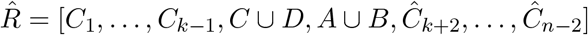

where for all *m* = *k* + 1, *…, n* − 2:

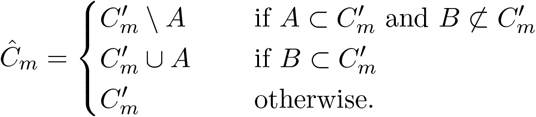

Using the fact that *B* ⊂ *C*_*m*_ if and only if 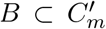, we can summarise the change of a cluster at rank *m > k* + 1 between *T* and 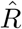 as follows:

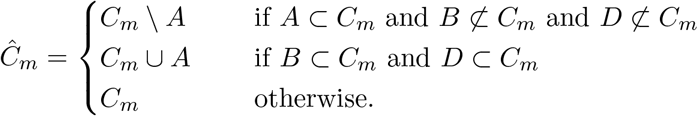

Since the cluster at rank *k* in *T* ′ is *A* ∪ *B*, it is *A* ⊂ *C*_*m*_ if and only if *B* ⊂ *C*_*m*_ for all *m > k*. Therefore, *Ĉ*_*m*_ = *C*_*m*_ for all *m > k* + 1. It follows that the cluster representation of 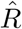 and *R* coincides, which gives us 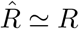, so *p*′ is a path from *T* to *R* with two HSPR moves (see Figure 9).
  2.2 If there is no edge between the nodes with rank *k* and *k* + 1 in *T*, then there is a cluster *E* in *T* so that *T* |_*A*_ is sibling of *T* |_*E*_ in *T* with *E*≠ *A, B, C, D* (see Figure 10). Then the cluster representation of *T* is:

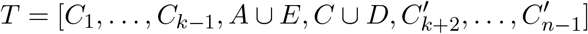

where for all *m* = *k* + 1, *…, n* − 2:

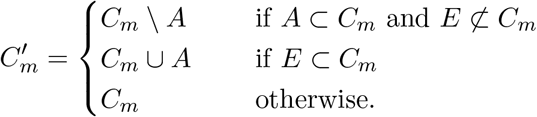 To create an alternative path *p*′, we first perform a rank move swapping the ranks of the nodes *k* and *k* + 1 in *T* to receive a tree *T* ″ with cluster representation:

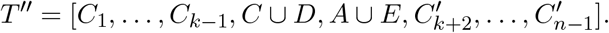

As second move on *p*′ we perform an HSPR move at rank *k* + 1, moving the subtree *T* ″|_*A*_ to become sibling of *T* ″|_*B*_, giving us the following tree 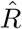:

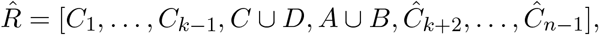

where clusters *Ĉ*_*m*_ with *m* ≥ *k* + 2 are defined as follows:

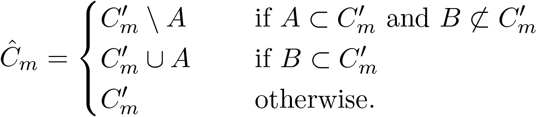

To show that *Ĉ*_*m*_ = *C*_*m*_ for all *m* = *k* + 2, *…, n* − 1, we distinguish whether *A* ⊂ *C*_*m*_ or *A* ⊄ *C*_*m*_. Note that *A* ⊂ *C*_*m*_ if and only if *B* ⊂ *C*_*m*_ and also *B* ⊂ *C*_*m*_ if and only if 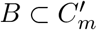. If *A* ⊂ *C*_*m*_ and *E* ⊄ *C*_*m*_, then 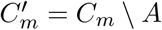 and since 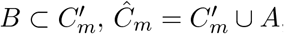, resulting in *Ĉ*_*m*_ = *C*_*m*_. If on the other hand *A* ⊂ *C*_*m*_ and *E* ⊂ *C*_*m*_, then 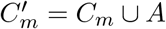 and with 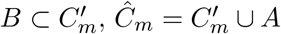, resulting in *Ĉ*_*m*_ = *C*_*m*_. If *A* ⊄ *C*_*m*_ and *E* ⊂ *C*_*m*_, then 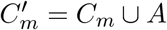 and since 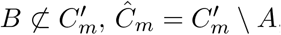, resulting in *Ĉ*_*m*_ = *C*_*m*_. If on the other hand *A* ⊄ *C*_*m*_ and *E⊄C*_*m*_, then 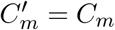 and since 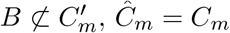, resulting in *Ĉ*_*m*_ = *C*_*m*_. So for every *m* = *k* + 2, *…, n* − 1, it is 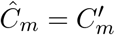, which implies 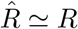, so *p*′ is a path from *T* to *R* with first a rank move and then an HSPR move.
3. *T* and *T* ′ are connected by an HSPR move at rank *k* + 1. As *T* |_*C*_ and *T* |_*D*_ are the subtrees that are children of node *k* + 1 in *T* ′, we can assume without loss of generality that *T* |_*C*_ is moved between *T* and *T* ′, otherwise we swap notation for *C* and *D*. We again further distinguish whether the node of rank *k* + 1 is parent of the node of rank *k* in *T* or not.
  3.1. Let there be an edge connecting nodes *k* + 1 and *k* in *T*. Remember that by our assumptions on *T* ′, parent_*T*′_ (*T* |_*A*_) = parent_*T*′_ (*T* |_*B*_) = *k*. Therefore, an HSPR move at rank *k* + 1 on *T* ′ that moves the subtree *T* |_*C*_ and creates a tree *T* containing an edge (*k* + 1, *k*) must move *T* |_*C*_ to become sibling of *T* |_*A∪B*_ (see Figure 11). Then *T* has the following cluster representation:

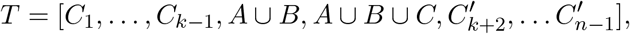

where

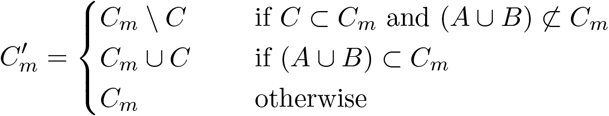

for all *m* = *k* + 2, *…, n* − 1. A path *p*′ can now be constructed by first performing an HSPR move at rank *k* on *T* moving *T* |_*A*_ to become sibling of *T* |_*C*_ in the resulting tree *T* ″:

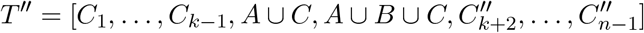

with

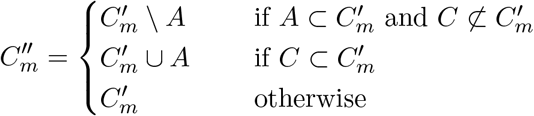

for all *m* = *k* + 2, *…, n* − 1. Because the node of rank *k* + 1 in *T* induced the cluster 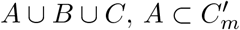 if and only if 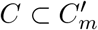 for all *m* = *k* + 2, *…, n* − 1. Therefore, 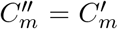 for all *m* = *k* + 2, *…, n* − 1. We then perform an HSPR move at rank *k* on *T* ″ that moves the subtree *T* ″|_*C*_ to become sibling of *T* ″|_*D*_, resulting in the following tree 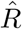:

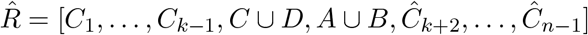

with

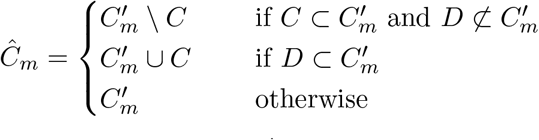

for all *m* = *k* + 2, *…, n* − 1. To show 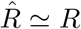, we use that *C* ⊂ *C*_*m*_ if and only if *D* ⊂ *C*_*m*_, and *A* ⊂ *C*_*m*_ if and only if *B* ⊂ *C*_*m*_ for all *m* = *k* + 2, *…, n* − 1, because the clusters induced by nodes of rank *k* and *k* + 1 in *T* ′ are *A* ∪ *B* and *C* ∪ *D*, respectively. Furthermore, we distinguish whether (*A* ∪ *B*) ⊂ *C*_*m*_, (*C* ∪ *D*) ⊂ *C*_*m*_, and (*A* ∪ *B* ∪ *C* ∪ *D*) ∩ *C*_*m*_ = ∅ If (*A* ∪ *B*) ⊂ *C*_*m*_ for some *m* = *k* + 2, *…, n* − 1, then 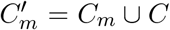. Then (i) 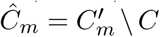 if 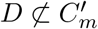 or (ii) 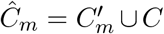 if 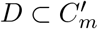. Note that if 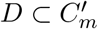, it must be *D* ⊂ *C*_*m*_, as *D* is not removed from any clusters between trees *T* ′ and *T*. And since *D* ⊂ *C*_*m*_ if and only if *C* ⊂ *C*_*m*_, we can see that 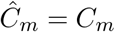 for both cases (i) and (ii). If (*C* ∪ *D*) ⊂ *C*_*m*_ for some *m* = *k* + 2, *…, n* − 1, then (i) 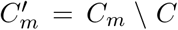 if (*A* ∪ *B*) ⊄ *C*_*m*_ or (ii) 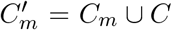 if (*A* ∪ *B*) ⊂ *C*_*m*_. Again, it is 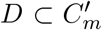 if and only if *D* ⊂ *C*_*m*_. Therefore, it must be 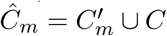 ∪ *C* in both cases (i) and (ii). With *C* ⊂ *C*_*m*_, it follows *Ĉ*_*m*_ = *C*_*m*_. If (*A* ∪ *B* ∪ *C* ∪ *D*) ∩ *C*_*m*_ = ∅, then *C*′*m* = *C*_*m*_ and 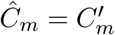, hence *Ĉ*_*m*_ = *C*_*m*_. We can conclude that 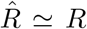, so *p*′ is a path from *T* to *R* consisting of two HSPR moves.
  3.2 There is no edge between the nodes of rank *k* and *k* + 1 in *T*. Let *E* be a cluster in *T* so that the subtree *T* |_*E*_ is sibling of *T* |_*C*_ in *T* with *E* ≠ *A, B, C, D*, i.e. *T* |_*C*_ is moved to become sibling of *T* |_*E*_ by the HSPR move between *T* and *T* (see Figure 12). The cluster notation for *T* is:

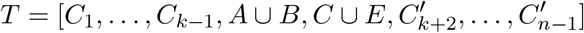

with

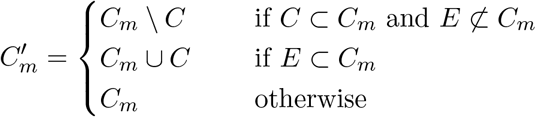

for all *m* = *k* + 2, *…, n* − 1. We construct a path *p*′ by first performing a rank move swapping the nodes *k* and *k* + 1 of *T*, resulting in the tree *T* ″:

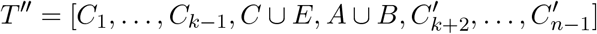

We can then preform an HSPR move on *T* ″, moving the subtree *T* ″|_*C*_ to become sibling of *T* ″|_*D*_, which gives us the tree 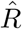:

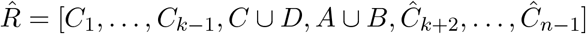

with

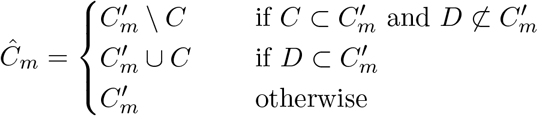

for all *m* = *k* + 2, *…, n* − 1. To show that *Ĉ*_*m*_ = *C*_*m*_ for all *m* = *k* + 2, *…, n* − 1, we distinguish whether *C* ⊂ *C*_*m*_ or *C* ?⊂ *C*_*m*_. Note that since *C* ∪ *D* is the cluster induced by node *k* in *T*, it is *C* ⊂ *C*_*m*_ if and only if *D* ⊂ *C*_*m*_ and since *D* is not removed from any clusters between *T* ′ and *T*, also *D* ⊂ *C*_*m*_ if and only if 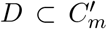for all *m* = *k* + 2, *…, n* − 1. If *C* ⊂ *C*_*m*_ and *E* ⊄ *C*_*m*_, then 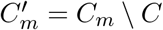and since 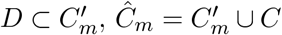, resulting in *Ĉ*_*m*_ = *C*_*m*_. If on the other hand *C* ⊂ *C*_*m*_ and *E* ⊂ *C*_*m*_, then 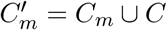 and with 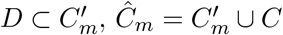, resulting in 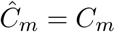. If *C* ⊄ *C*_*m*_ and *E* ⊂ *C*_*m*_, then 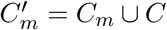 and since 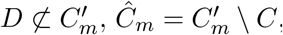 resulting in *Ĉ*_*m*_ = *C*_*m*_. If on the other side *C* ⊄ *C*_*m*_ and *E* ⊄ *C*_*m*_, then 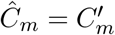 and 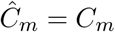, resulting in *Ĉ*_*m*_ = *C*_*m*_. In any case, it is *Ĉ*_*m*_ = *C*_*m*_ for all *m* = *k* + 2, *…, n* − 1 and therefore 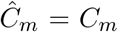, which implies that *p* is a path from *T* to *R* that consists of a rank move followed by an HSPR move.

**Figure 10.**
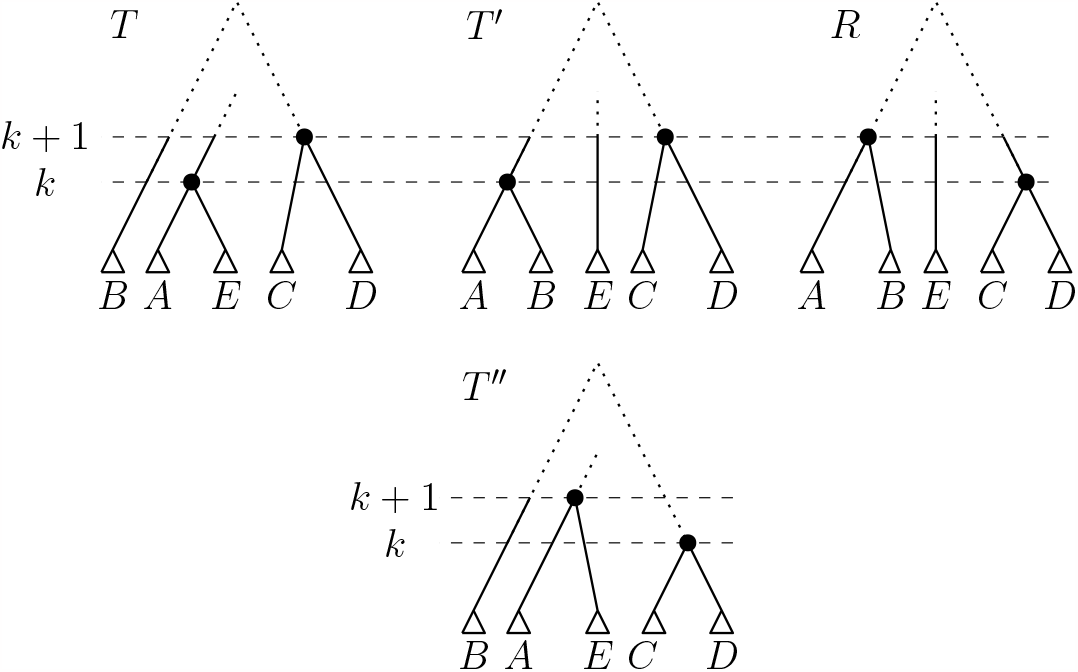
Trees *T, T* ′, and *R* on *p* at the top, and alternative path *p*′ from *T* to *R* via *T* ″ at the bottom if *T* and *T* ′ are connected by an HSPR move at rank *k* and the nodes of rank *k* and *k* + 1 in *T* are not connected by an edge. The dotted parts of the trees might contain further nodes and leaves.

**Figure 11.**
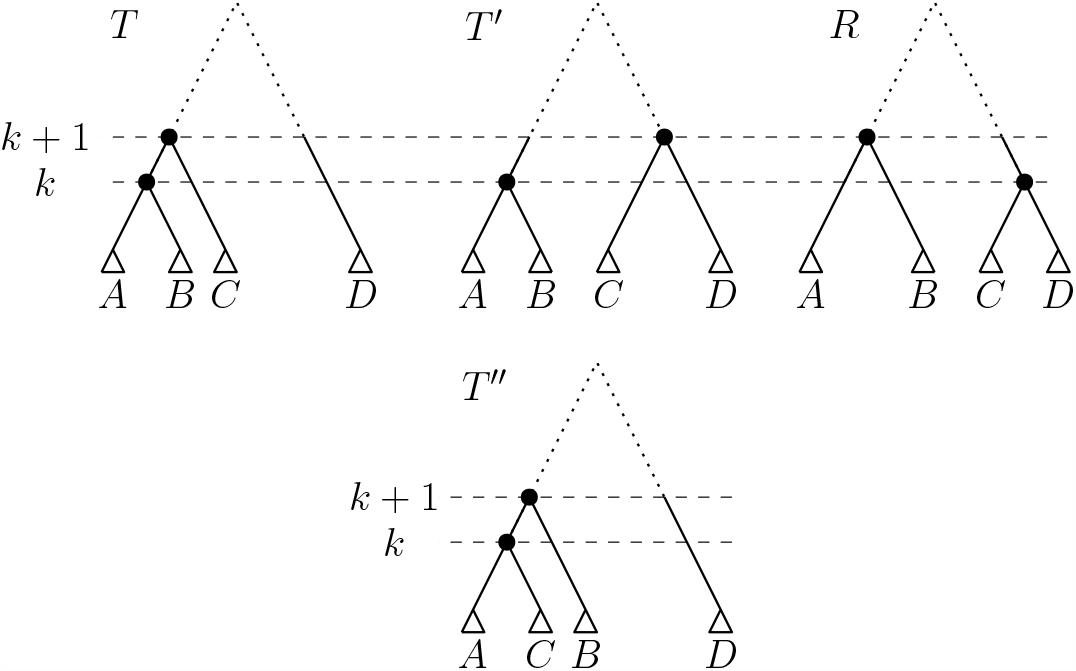
Trees *T, T* ′, and *R* on *p*, and alternative path *p*′ from *T* to *R* via *T* ″ at the bottom if *T* and *T* ′ are connected by an HSPR move at rank *k* + 1 and the nodes of rank *k* and *k* + 1 in *T* are connected by an edge, The dotted parts of the trees might contain further nodes and leaves.

**Figure 12.**
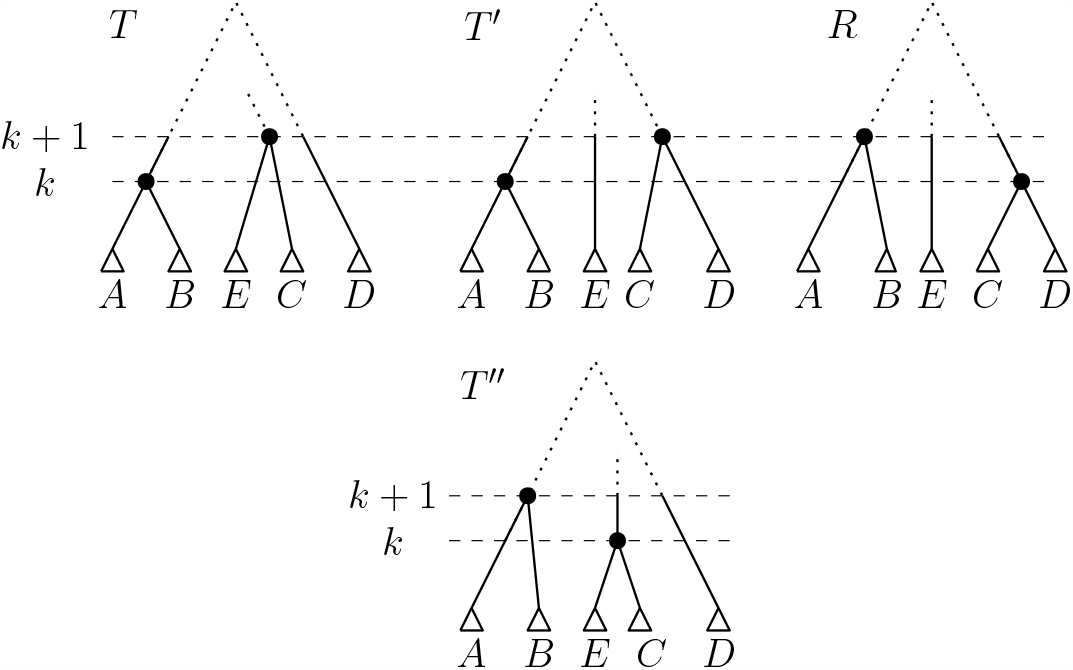
Trees *T, T* ′, and *R* on *p* at the top, and alternative path *p*′ from *T* to *R* via *T* ″ at the bottom if *T* and *T* ′ are connected by an HSPR move at rank *k* and the nodes of rank *k* and *k* + 1 in *T* are not connected by an edge. The dotted parts of the trees might contain further nodes and leaves.

In all cases above, we found an alternative path *p*′ to the path *p* so that *p*′ consists of either two HSPR moves or a rank move followed by an HSPR move, which concludes the proof of this lemma.

Using Lemma 5, we can now prove that we can change the order of moves on an RSPR path so that in the beginning we have a sequence of rank moves, followed by a sequence of HSPR moves.

#### Theorem 10

*In* RSPR *there is always a shortest path between two trees that has a sequence of rank moves (if there are any) at the beginning followed by only* HSPR *moves*.

*Proof*. To prove the theorem we assume that there is a shortest path *p* = [*T*_0_, *T*_1_, *…, T*_*d*_] between trees *T*_0_ and *T*_*d*_ that has at least one rank move preceded by an HSPR move. Let *T*_*i*−1_ and *T*_*i*_ for some 1 *< i < d* − 1 be connected by the first HSPR move on *p* that has a rank move following it, i.e. *T*_*i*_ and *T*_*i*+1_ are connected by a rank move. By Lemma 5, we can replace *T*_*i*_ by a tree 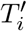 so that there is a rank move or HSPR move between *T*_*i*−1_ and 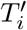 and an HSPR move between 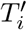 and *R*. When iteratively applying this procedure to all rank moves that are preceded by an HSPR move on *p*, we receive a path from *T* to *R* that has all rank moves at the beginning of the path, followed by a sequence of only HSPR moves.

Theorem 10 implies that any (not necessarily shortest) path between two trees *T* and *R* can be converted into a *T* -*R*-path of the same length that has all rank moves bundled at the beginning of the path, followed by HSPR moves. We can also shuffle moves on an RSPR path in the opposite way so that HSPR moves happen first and rank moves last. Additionally, we can infer the following from Theorem 10.

#### Corollary 5

*Let T*_*u*_ *and R*_*u*_ *be rooted (unranked) trees. Then there are ranked trees T* = (*T*_*u*_, rank_*T*_) *and R* = (*R*_*u*_, rank_*R*_) *so that d*_RSPR_(*T, R*) = *d*_HSPR_(*T, R*).

*Proof*. Let 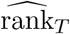 be an arbitrary rank function of *T*_*u*_, i.e. 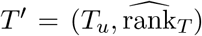is a ranked tree, and let rank_*R*_ be a rank function on *R*_*u*_ so that *R* = (*R*_*u*_, rank_*R*_) is a ranked tree. By Theorem 10, we can find a shortest path *p* from *T* ′ to *R* in RSPR that has all rank moves bundled in the beginning of the path. Let *T* be the ranked tree on *p* after this sequence of rank moves, i.e. the remainder of *p* between *T* and *R* consists of HSPR moves only. Since there is a path from *T* ′ to *T* consisting of rank moves only, the unranked versions of *T* ′ and *T* are identical, i.e. *T* = (*T*_*u*_, rank_*T*_) for some rank function rank_*T*_ on *T*_*u*_. Furthermore, as *p* is a shortest path in RSPR, the part of *p* between *T* and *R* is a shortest path between those trees in RSPR. And because this part of *p* consists of HSPR moves only, it must also be a shortest path in HSPR. We conclude that there are ranked trees *T* = (*T*_*u*_, rank_*T*_) and *R* = (*R*_*u*_, rank_*R*_) so that *d*_RSPR_(*T, R*) = *d*_HSPR_(*T, R*).

#### Corollary 6

*If there is a shortest path between trees T and R in* RSPR *that contains a caterpillar tree, then there is a shortest path from T to R that consists of* HSPR *moves only and d*_HSPR_(*T, R*) = *d*_RSPR_(*T, R*).

*Proof*. Let *T* and *R* be trees connected by a shortest path *p* that contains a caterpillar tree *T*_*c*_. The sequence of trees on *p* between *T* and *T*_*c*_ is a shortest path between these trees, and its reversed order a shortest path from *T*_*c*_ to *T*. Applying Theorem 10 to this path from *T*_*c*_ to *T* gives a shortest path where all rank moves are at the beginning of the path. Since *T*_*c*_ is a caterpillar tree, there cannot be rank moves on *T*_*c*_ which implies that there is a shortest paths from *T* to *T*_*c*_ that consists of HSPR moves only.

By the same argument, the restriction of *p* from *T*_*c*_ to *R* can be transformed into a path of the same length that consists of only HSPR moves.

We can concatenate the shortest path from *T* to *T*_*c*_ with only HSPR moves and the one from *T*_*c*_ and *R* with only HSPR moves and receive a path from *T* to *R* that has the same length as *p*. This path is hence a shortest path and consists of HSPR moves only.

## 5 Adding leaves

In this section we consider how adding a leaf to two trees can change their distance. Especially when considering data sets of ongoing evolutionary processes, like virus transmissions, it would be ideal to re-use the already inferred tree, and some methods to do this already exist [6, 13, 15, 17]. For analyses where new leaves are added to an already existing tree, it is important to understand how adding a leaf can change a tree. It is also of interest to know how the distance between two trees changes when a new leaf is added. One would naturally assume that the addition of a leaf to two trees increases their distance. For HSPR and RSPR, however, we find that adding one leaf can decrease the distance between two trees linearly in *n* (Theorem 11). We will see that this is the case when two trees are similar and the new leaf is added at different heights in the tree. Such leaves with varying positions in a tree are often referred to as “rogue taxa” [1], and our results here show that the existence of these can have a big impact on the distance between two trees under RSPR and HSPR.

Before we describe how adding a leaf results in a decrease in distance, we need the following observation.

### Lemma 6

*The caterpillar trees*

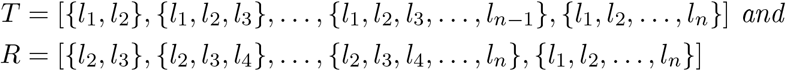

*have* HSPR *and* RSPR *distance greater than or equal to* 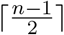.

The trees *T* and *R* of Lemma 6 are displayed in the top row of Figure 13. Whether the bound for the distance of *T* and *R* in Lemma 6 is sharp remains an open question.

*Proof*. The parents of *n* − 1 leaves *l*_1_, *l*_3_, *l*_4_, *…, l*_*n*_, i.e. all leaves except *l*_2_, have different ranks in *T* and *R*. By Lemma 1 it follows 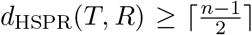. And as the HSPR and RSPR distance between caterpillar trees coincides (Corollary 6), 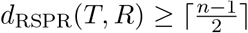.

**Figure 13.**
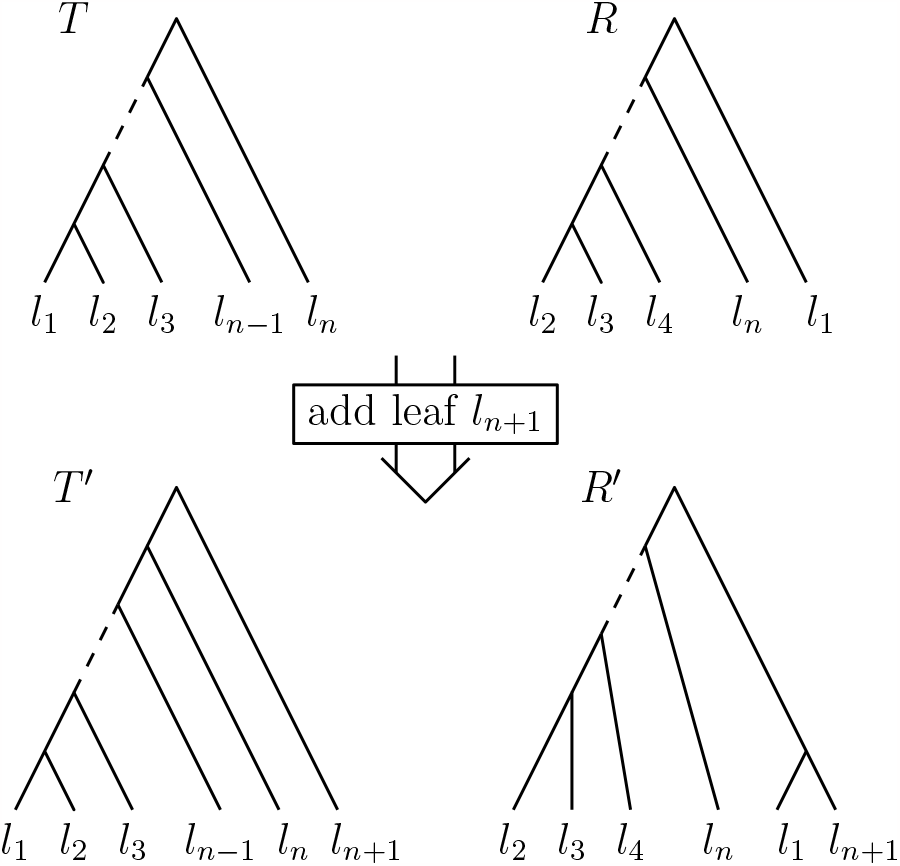
Top: trees *T* and *R* with HSPR distance greater than or equal to ^*n*−1^ (Lemma 6). Bottom: Adding leaf *l*_*n*+1_ to both *T* and *R* results in trees *T* ′ and *R*′ that are connected by one HSPR move that moves *l*_1_.

Note that Lemma 6 implies that the diameter of RSPR is greater than or equal to 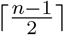, which is the same boundary we already found for HSPR in Theorem 6. We are now ready to prove the main theorem of this section.

### Theorem 11

*Adding a leaf to two trees can decrease their* HSPR *distance by* 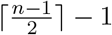. *Proof*. Let *T* and *R* be the trees of Lemma 6. We add a leaf *l*_*n*+1_ to these trees *T* and *R* as follows (see Figure 13): A new root is added to *T* so that the resulting tree *T* ′ has *l*_*n*+1_ and the old root of *T* as children. In *R* we attach *l*_*n*+1_ as sibling of *l*_1_ so that their parent has rank one, and the ranks of all other internal nodes of *R* are increased by one, giving us a tree *R*′ on *n* + 1 leaves. The cluster representations of *T* ′ and *R*′ are:

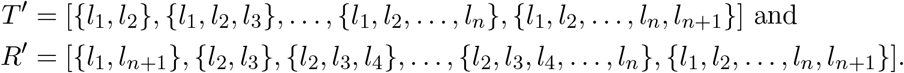

Let *C*_*i*_ and 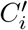 be the cluster induced by the node *i* in *T* ′ and *R*′, respectively, for all *i* = 1, *…, n* − 1. We can then describe the difference between the cluster representations of *T* ′ and *R*′ as follows: 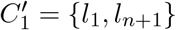 and for all *m* = 2, *…, n*:

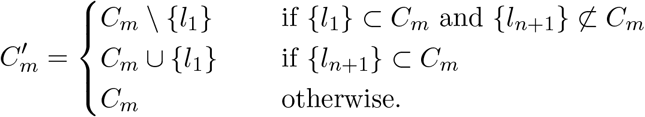

By Theorem 3, *T* ′ and *R*′ are connected by an HSPR move.

Therefore, adding one leaf to *T* and *R* changes the distance from 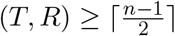 (Lemma 6) to *d*_HSPR_(*T*′, *R*′) = 1, which means that their distance decreases by at least 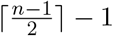.

In Theorem 12 we will see that adding a leaf can only increase the HSPR distance if the unranked versions of the given trees are connected by a rooted (unranked) SPR move.

### Theorem 12

*Let T* = (*T*_*u*_, rank_*R*_) *and R* = (*R*_*u*_, rank_*R*_) *be trees with* HSPR *distance d*_HSPR_(*T, R*) = *d >* 1, *and let T* ′ *and R*′ *be trees resulting from adding a leaf x to T and R, respectively, so that d*_HSPR_(*T* ′, *R*′) = 1.

*If T*_*u*_ *and R*_*u*_ *are the unranked versions of T and R, respectively, then the rooted (unranked)* SPR *distance of T*_*u*_ *and R*_*u*_ *is one: d*_SPR_(*T*_*u*_, *R*_*u*_) = 1.

*Proof*. Since *T* ′ and *R*′ have HSPR distance one, there is a subtree *T*| _*i*_ present in *T* ′ and *R*′ that is moved between them. It follows directly from the definitions of SPR and HSPR moves that if *T* ′ and *R*′ are connected by an HSPR move, their unranked versions 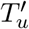 and 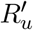 are connected by an SPR move, too, and the subtree moving between 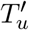 and 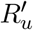 is the unranked version 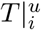 of *T* |_*i*_.

To show *d*_SPR_(*T*_*u*_, *R*_*u*_) = 1, we compare the distance of *T*_*u*_ and *R*_*u*_ with that of 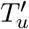 and 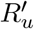 and analyse how removing leaf *x* from 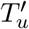 and 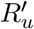 to obtain *T*_*u*_ and *R*_*u*_ affects their distance. We know that 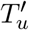 and 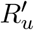 differ only in the position of subtree 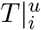. We consider following cases, depending on the position of the leaf *x* relative to 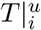:

i. If 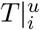 has *x* as its only leaf, then *T*_*u*_ ≃ *R*_*u*_, contradicting the assumptions of the theorem.
ii. If *x* is not in 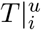, then *T*_*u*_ and *R*_*u*_ differ only by the position of 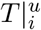.
iii. If *x* is not the only leaf in the leaf set of 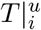, then *T*_*u*_ and *R*_*u*_ differ only by the position of 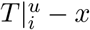.

While case (i) is not possible under our assumptions, case (ii) and (iii) both imply that *T*_*u*_ and *R*_*u*_ are connected by one SPR move. Therefore, we conclude that *d*_SPR_(*T*_*u*_, *R*_*u*_) = 1.

## 6 Discussion

Many tree inference methods use SPR moves for tree proposals, and extensive research made the SPR treespace usable for analysing tree inference methods and their output. There have however not been any comparable studies for phylogenetic time trees, even though time trees are inferred from sequence data in many applications and software packages like BEAST2 [7] for time tree inference are extremely popular. One approach to considering SPR moves for time trees is that by Song [32], where SPR moves are allowed to re-attach subtrees at the same height or closer to the root than its previous attachments. With this paper, we introduce two ranked SPR treespaces, RSPR and HSPR, that are motivated by the variations of SPR moves that are actually used as tree proposals in practice and have some properties in common with rooted (unranked) SPR. We show that all of these treespaces are connected, have neighbourhood sizes quadratic in the number of leaves *n*, and have a diameter linear in *n*. These properties have already been proven to be useful for SPR, which provides a good range of different trees for tree proposals [36]. Adding ranks to SPR provides an even more biologically relevant distance measure, as it ensures that the times of nodes in subtrees cannot change by one HSPR move, which particularly makes sense when using the number of SPR moves as a proxy for the number of reticulation events such as hybridisation, recombination, or horizontal gene transfer. Furthermore, HSPR moves provide a wider range of trees at close distance than for example RNNI moves, which is especially useful for tree proposals. These observations demonstrate that studying ranked SPR treespaces may provide insights to better understanding phylogenetic methods for time trees.

We also find some interesting differences between ranked and unranked SPR spaces. One of those is the absence of the (weak and strong) cluster property in ranked SPR. This suggests that the work on Maximum Agreement Forests (MAFs) cannot be transferred to ranked SPR spaces (see [37] for a discussion on MAF-like problems). Since MAFs are essential in the proofs of 𝒩 𝒫 -hardness for classic SPR, a different strategy is needed to prove the complexity of computing distances in RSPR and HSPR. Note that it is currently not known whether this problem is 𝒩 𝒫-hard in the ranked SPR treespaces.

We did however obtain some results on properties for shortest paths in HSPR and RSPR. The order of moves on shortest paths in RSPR can be changed so that first rank moves and then HSPR moves are performed, which suggests that the complexity of computing distances in the two spaces is the same. Furthermore, we found that there is a shortest path on which the ranks of HSPR moves increases monotonically. Leveraging these results in ranked SPR spaces might lead to developing an algorithm for computing or approximating distances in these treespaces. Another characteristic of ranked SPR spaces that makes them stand out from known tree rearrangement based treespaces is that adding one leaf to two trees can decrease their distance linearly in *n*. This is very interesting behaviour, as it has not been observed before in any of the known tree rearrangement based treespaces, and seems to be related to the presence of rogue leaves. It is worthwhile investigating the influence of this effect on tree inference algorithms, especially when considering algorithms that aim to add new sequence data to existing phylogenies (online algorithms).

This paper provides the definition and first analysis of spaces of ranked trees to enable studying time tree inference methods in the same way as untimed tree inference methods. One important open question is that of the complexity of computing distances in RSPR and HSPR treespace. A first step could be to establish that the complexity of computing distances is the same for these two ranked treespaces. Furthermore, the exact diameter of HSPR and RSPR space is still to be determined. We can use our computations [8] to show that the diameter of HSPR follows the formula 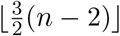 for *n* ≤ 7, but whether this is true for any *n* remains an open question. Another very important next step for research on ranked SPR is developing algorithms, ideally fixed-parameter tractable ones, to calculate or approximate distances. This would facilitate leveraging our newly defined treespaces to allow analysing BEAST2 [7] output as it has been done for MrBayes output with unranked SPR [36, 38].

## Footnotes

1 https://beast.community/xml_reference#ltfixednodeheightsubtreepruneregraftgt-element

## Notes

This work was partially supported by Royal Society Te Apārangi through a Rutherford Discovery Fellowship (UOC1702) and a Marsden grant (21-UOC-057), and by Ministry of Business, Innovation, and Employment of New Zealand through an Endeavour Smart Ideas grant (UOOX1912) and a Data Science Programmes grant (UOAX1932).

### Competing Interest Statement

The authors have declared no competing interest.

### Summary of Updates

We introduced new notation in Section 2 (Preliminaries) to increase clarity of proofs throughout the manuscript.

